# Peptide-mimetic treatment of *Pseudomonas aeruginosa* in a mouse model of respiratory infection

**DOI:** 10.1101/2023.10.30.564794

**Authors:** Madeleine G. Moule, Aaron B. Benjamin, Melanie L. Buger, Claudine Herlan, Maxim Lebedev, Jennifer S. Lin, Kent J. Koster, Neha Wavare, Leslie G. Adams, Stefan Bräse, Annelise E. Barron, Jeffrey D. Cirillo

**Author notes:** These authors had equal contributions to this manuscript (Madeleine G. Moule, Aaron B. Benjamin). Co-corresponding authors: Annelise E. Barron Jeffrey D. Cirillo.

## Abstract

The rise of drug resistance has become a global crisis, with >1 million deaths due to resistant bacterial infections each year. *Pseudomonas aeruginosa,* in particular, remains a serious problem with limited solutions due to complex resistance mechanisms that now lead to more than 32,000 multidrug-resistant (MDR) infections and over 2,000 deaths annually. While the emergence of resistant bacteria has become concerningly common, identification of useful new drug classes has been limited over the past 40+ years. We found that a potential novel therapeutic, the peptide-mimetic TM5, is effective at killing *P. aeruginosa* and displays sufficiently low toxicity for mammalian cells to allow for use in treatment of infections. Interestingly, TM5 kills *P. aeruginosa* more rapidly than traditional antibiotics, within 30-60 minutes *in vitro*, and is effective against a range of clinical isolates. *In vivo*, TM5 significantly reduced bacterial load in the lungs within 24 hours compared to untreated mice and demonstrated few adverse effects. Taken together, these observations suggest that TM5 shows promise as an alternative therapy for MDR *P. aeruginosa* respiratory infections.

## Introduction

The rising prevalence of antimicrobial resistance (AMR) has resulted in a resurgence of bacterial diseases that would otherwise have been successfully treated with antibiotics. In 2019, the WHO reported that at least 700,000 people die each year due to drug-resistant diseases and predicted that this number could rise to 10 million deaths per year by 2050^1^. A surveillance report by the European Center for Disease Control (eCDC) using data from 2021 reported that 6 of 44 countries in Europe had antimicrobial resistance rates over 50% while only 2 of 44 European countries had rates less than 5% ^2^. The American Center for Disease Control (CDC) outlined the biggest AMR threats to the United States in a 2019 report, one of which is the facultative aerobe *Pseudomonas aeruginosa*^3^.

*P. aeruginosa* is a ubiquitous, opportunistic pathogen which utilizes over 100 different organic molecules for energy acquisition via carbon or as a prototroph ^4^. One significant problem with treatment of *P. aeruginosa* infections is the ability of the bacteria to develop antibiotic resistance quickly via both horizontal gene transfer and mutations which lead to up-regulation of β-lactamases and efflux pumps ^4, 5^. The most severe *Pseudomonas* infections include respiratory infections in patients with cystic fibrosis and systemic bloodstream infections that have disseminated from burn wounds or pneumonia ^5, 6^. In the United States alone, it was estimated that over 30,000 multidrug-resistant cases were seen in 2019, with estimated healthcare associated costs of $750 million in 2017 ^3^. Due to this increase in resistance, there is an urgent need to develop new antimicrobials against *P. aeruginosa* which can bypass traditional resistance mechanisms.

The current frontline treatments for infections with *P. aeruginosa* encompass a variety of different antibiotic classes, including aminoglycosides, carbapenems, cephalosporins, fluoroquinolones, penicillins with β-lactamases, monobactams, and Fosfomycin with polymyxins as a last resort ^5, 7, 8^. Despite this diversity of options, misuse of antibiotics and noncompliance has increased the amount of drug resistance observed in clinical infections and resulted in the emergence of multidrug resistant (MDR) strains. Additionally, since the golden era of antibiotic discovery ended in the 1960s, traditional approaches to drug discovery have yielded fewer new classes of antimicrobials each subsequent year ^9–11^. While new antibiotic development has been stuck at a snail’s pace, the bacterial world has continued to evolve and develop resistances at an alarming rate, resulting in a wide range of mechanisms through which bacteria can overcome antimicrobial burden. Examples include, but are not limited to, efflux pumps, modification of antibiotics, changes in the membrane to prevent drug penetration, and target modification ^9, 12^. To bypass these methods of resistance, alternative strategies need to be explored for drug development, such as utilizing natural host-based immunity via biomimetics.

One family of promising antibiotic alternatives are antimicrobial peptides (AMPs). AMPs exhibit an extensive spectrum of applications and a lower risk of resistance development compared to traditional antibiotics, making them promising candidates as next-generation antibiotics ^13^. AMPs, also called host defense peptides, are produced by nearly all organisms from prokaryotes to humans as products of the innate immune system. Most AMPs are short cationic amphiphilic peptides and display activity against a broad range of pathogens including bacteria, viruses, and fungi ^14^. They have also demonstrated anticancer properties ^15^, have been seen as chemoattractants ^16^, and can modulate the inflammatory response ^17–19^. In humans, endogenous AMPs are grouped into three families based on their structural homology motifs: defensins, cathelicidins and histatins ^20–24^. In humans, there is only one known cathelicidin, the host defense peptide LL-37 ^25, 26^. LL-37 is involved in a variety of functions, including both direct microbicidal activities and immunomodulatory functions ^18, 19, 27–33^. Unfortunately, LL-37 is not able to eliminate all bacterial infections by itself, and resistance against some natural antimicrobial peptides has begun to increase ^34^. Additionally, their low bioavailability and susceptibility to degradation by proteases *in vivo* reduces their therapeutic potential. Despite this, their efficacy against both gram-positive and gram-negative bacteria, including antibiotic resistant strains, makes AMPs potential scaffolds to for the design of mimics capable of treating infections ^35^.

Since AMPs are easily manipulated and their structures are so tightly connected to their efficacy, creating peptidomimetics provides an untapped reservoir of antimicrobial therapeutic solutions. We have previously developed a class of AMP derivates known as peptoids, or *N*-substituted glycine oligomer peptidomimetics, based off LL-37 ^36–39^. Peptoids differ from peptides in that the side chains are linked to the amide in the backbone instead of the α-carbon. This change in structure removes hydrogen bonding from the backbone resulting in resistance to proteolysis. Here, we focus on the therapeutic potential of the peptoid TM5 against *P. aeruginosa*. We have shown that TM5 is a promising candidate for treatment of *P. aeruginosa* infections due to its potent antimicrobial activity against this organism, improved stability compared to the parental AMP, and low toxicity to the host.

## Results

### Peptoid TM5 is an effective antimicrobial against *P. aeruginosa* Xen41

AMPs have previously demonstrated potent antimicrobial activity *in vitro* against a wide range of gram-negative and gram-positive bacteria, including *P. aeruginosa* ^40–42^. To evaluate efficacy of our peptoids against *P. aeruginosa*, we first determined the minimum inhibitory concentration (MIC) and minimum bactericidal concentration (MBC) of TM5 against the bioluminescent strain Xen41. Xen41 is a strain derived from the parental strain *P. aeruginosa* PAO1 that contains an integrated LuxABCDE operon cloned from *Photorhabdus luminescens* that provides strong constitutive bioluminescence suitable for *in vivo* imaging studies ^43, 44^. To determine the MIC, overnight cultures of *P. aeruginosa* were treated with a range of 0 – 64 µg/mL of several possible treatments including, TM1, TM5, TM6, ceftazidime, ciprofloxacin, kanamycin, and meropenem. Each sample was tested in triplicate, and OD_600_ and luminescence were measured following incubation at 37°C for 4 hours. The MIC_90_s for TM1 and TM5 were determined to be 4-8 and 4 µg/mL respectively, while the MIC_90_ for TM6 was estimated at 32 µg/mL (Figure 1a). MICs were also determined based off 50% reduction of luminescence (MIC_50_). MIC_50_s for each peptoid was half that of the MIC_90_ (Figure 1a). At concentrations of 1 µg/mL and higher, all three peptoids showed significant reductions in relative luminescence (Figure 1a).

**Figure 1:**
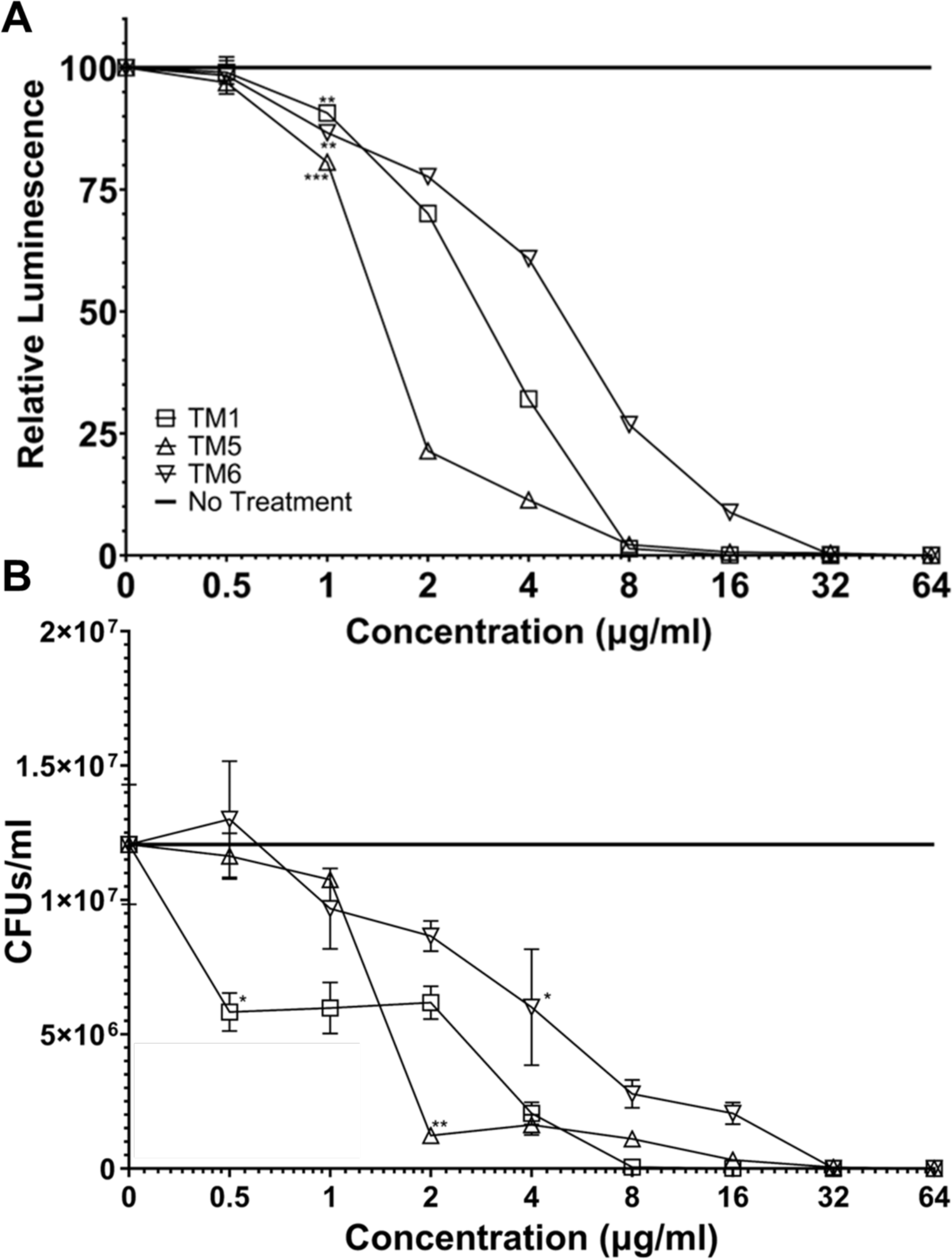
Relative luminescence (A) and CFUs/ml (B) for *P. aeruginosa* Xen41 with 2-fold dilutions of TM1, TM5, and TM6. (A) TM1, TM5, and TM6 were all significantly different from the no treatment control with respect to their relative luminescence at concentrations of 1 μg/ml and higher. (B) TM1 was significantly different from the no treatment control for CFUs/ml at concentrations of 0.5 μg/ml and above, while TM5 and TM6 were significantly different at 2 and 4 μg/ml respectively. Data points are represented as means using four replicates. Error is shown in ± standard deviation (SD). Statistics were performed using 2-way ANOVA, comparing antimicrobial to no treatment control. *P* values are: <0.001 = ***, between 0.001 and 0.01 = **, and between 0.01 and 0.05 = *.

To determine the MBC, an indicator of bacterial killing rather than growth inhibition, each well was diluted and spotted onto LB agar and bacterial colony forming units (CFU) were enumerated. The MBC of TM1 and TM6 were 8 and 32 µg/mL respectively, (Figure 1b). For TM5, the MBC was determined to be between 2-16 µg/mL, as the CFUs/mL hovered at the MBC threshold of 95% killing at all concentrations within that range (Figure 1b). TM1 showed a significant decrease in CFUs at concentrations higher than 0.5 µg/mL, TM5 at concentrations higher than 2 µg/mL, and TM6 at concentrations higher than 4 µg/mL. All findings pertaining to MIC and MBC for the peptoids are summarized in Table 1.

**Table 1.** MICs and MBC of peptoids against *P. aeruginosa* Xen41.

| **Peptoid** | **Sequence** | **MBC** | **MIC<sub>50</sub>** | **MIC<sub>90</sub>** |
| --- | --- | --- | --- | --- |
| TM1 | H-(NLys-Nspe-Nspe)<sub>4</sub>-NH<sub>2</sub> | 8 µg/ml | 2-4 µg/ml | 4-8 µg/ml |
| TM5 | H-Ntridec-NLys-Nspe-Nspe-NLys-NH<sub>2</sub> | 2-16 µg/ml | 1-2 µg/ml | 4 µg/ml |
| TM6 | H-(NLys-Nspe-Nspe)<sub>3</sub>-NLys-Nspe-NH<sub>2</sub> | 32 µg/ml | 4-8 µg/ml | 16 µg/ml |

As a control for comparative efficacy against *P. aeruginosa* Xen41, the antibiotics ceftazidime, ciprofloxacin, kanamycin, and meropenem were all tested in the same manner. Ciprofloxacin showed the lowest MIC_90_, MIC_50_, and MBC, all of which were <0.5 µg/mL (Figure 2). Interestingly, ceftazidime gave MICs of 1 µg/mL, but an MBC of 2-4 µg/mL (Figure 2). Kanamycin showed consistent MIC_90_ and MBC of >64 µg/mL, showing 50% luminescence reduction between 32 and 64 µg/mL (Figure 2). Meropenem had an MIC_90_ of 1 µg/mL (Figure 2a) and an MBC of 0.5 µg/mL (Figure 2b). Ciprofloxacin and meropenem both showed a significant decrease in luminescence and CFUs at concentrations of 0.5 µg/mL and higher, while ceftazidime showed a significant reduction of luminescence at all concentrations except 2 µg/mL and for CFUs at all concentrations (Figure 2). Kanamycin showed a significant decrease in luminescence for all concentrations above 1 µg/mL, but only showed a significant decrease of CFUs at concentrations higher than 4 µg/mL (Figure 2). All findings pertaining to MIC and MBC for the traditionally used antibiotics are summarized in Table 2.

**Figure 2:**
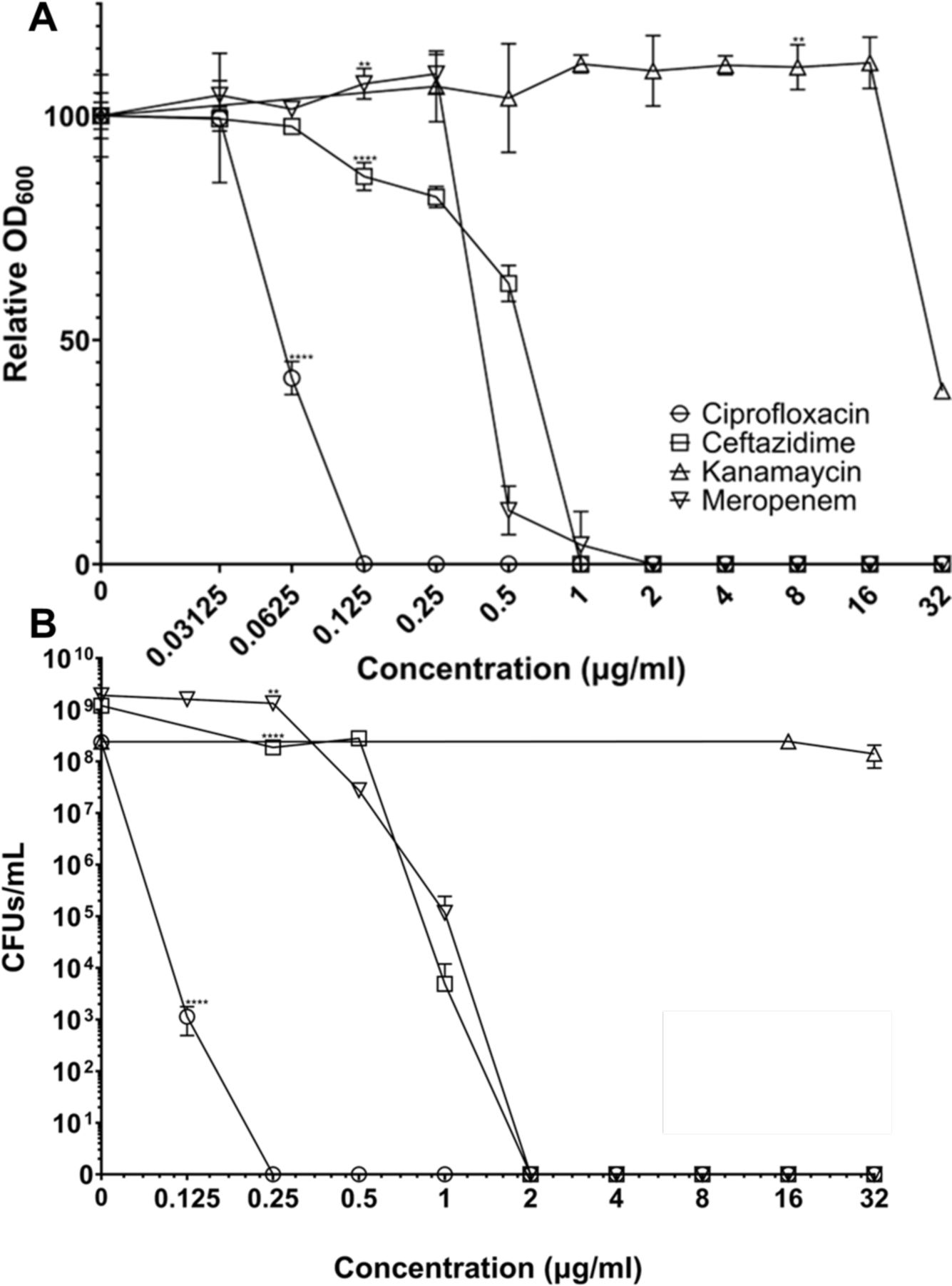
Relative OD_600_ (A) and CFUs/ml (B) for *P. aeruginosa* Xen41 with 2-fold dilutions of ciprofloxacin, ceftazidime, kanamycin, and meropenem. (A) Ciprofloxacin was significantly different above concentrations of 0.0625 μg/ml and above. Ceftazidime and meropenem were significantly different at concentrations above 0.125 μg/ml, while kanamycin was significantly different at concentrations of 8 μg/ml and above. (B) Ciprofloxacin showed a significant decrease compared to the no treatment control for CFUs/ml at all concentrations, while ceftazidime and meropenem were significantly different at 0.25 μg/ml and above. Kanamycin didn’t show any significant difference at any concentrations. Data points are represented as means using four replicates. Error is shown in ± standard deviation (SD). Statistics were performed using 2-way ANOVA, comparing antimicrobial to no treatment control. *P* values are: <0.0001 = ****, between 0.0001 and 0.001 = ***, between 0.001 and 0.01 = **, and between 0.01 and 0.05 = *.

**Table 2.**
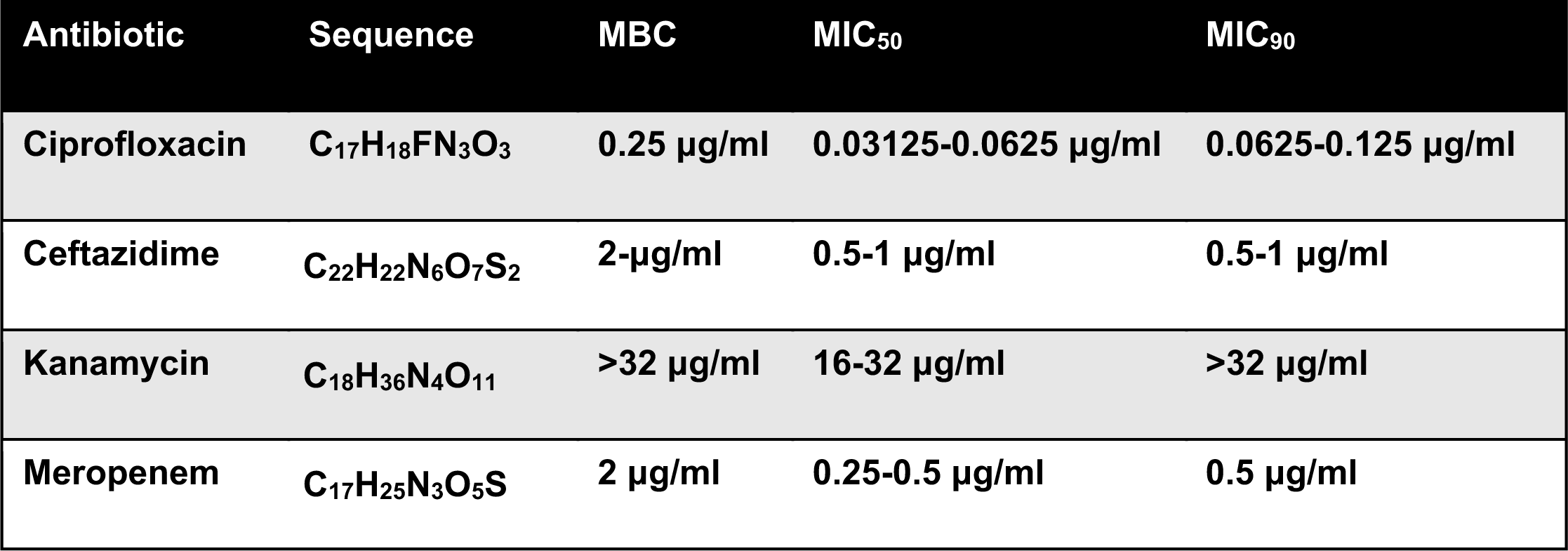
MICs and MBC of traditional antibiotics against *P. aeruginosa* Xen41.

### TM5 demonstrates consistent efficacy against clinical isolates of *P. aeruginosa*

*P. aeruginosa* PAO1, the parental strain from which Xen41 was derived, was originally a clinical isolate derived from a wound infection in an Australian patient in 1954. However, due to the genetic plasticity of *P. aeruginosa,* it is well documented that continuous propagation has resulted in laboratory strains demonstrating significant differences in virulence and antibiotic resistance profiles compared to direct clinical isolates. ^45, 46^ To determine if our peptoids retain the ability to effectively kill clinical strains from diverse sources with different AMR profiles, we tested our most promising candidate, TM5, against clinical strains from the Kolter collection^47, 48^ We compared the MIC and MBC of a second laboratory strain from this collection that demonstrates increased virulence (PA14) against strains isolated from infections of a trachea (PAZK69), wound (PAZK2019), and from a cystic fibrosis lung infection (PAZK2870). Although these strains have a variety of antibiotic susceptibility profiles, every strain demonstrated consistent susceptibility to peptoid TM5, with all strains tested showing a MIC of 8 ug/mL, and MBCs of 16 µg/mL (Table 3, Table 4)

**Table 3.** MICs of TM5 and traditional antibiotics against clinical strains of *P. aeruginosa*.

| Peptoid/<br>Antibiotic | Sequence | MIC |  |  |  |  |
| --- | --- | --- | --- | --- | --- | --- |
|  |  | PA14 | PAZK2870<br>YFP | PAZK2019<br>YFP | PAZK69<br>YFP | PAZK3095<br>YFP |
| TM5 | H-Ntridec-NLys-Nspe-Nspe-NLys-NH<sub>2</sub> | 8 µg/ml | 8 µg/ml | 8 µg/ml | 8 µg/ml | - |
| Ciprofloxacin | C<sub>17</sub>H<sub>18</sub>FN<sub>3</sub>O<sub>3</sub> | 0.125<br>µg/ml | 0.125 µg/ml | 0.125 µg/ml | 0.06<br>µg/ml | 0.25 µg/ml |
| Ceftazidime | C<sub>22</sub>H<sub>22</sub>N<sub>6</sub>O<sub>7</sub>S<sub>2</sub> | 2 µg/ml | 2 µg/ml | 2 µg/ml | 1 µg/ml | 1 µg/ml |
| Kanamycin | C<sub>18</sub>H<sub>36</sub>N<sub>4</sub>O<sub>11</sub> | 32 µg/ml | 64 µg/ml | 64 µg/ml | 32 µg/ml | 64 µg/ml |
| Meropenem | C<sub>17</sub>H<sub>25</sub>N<sub>3</sub>O<sub>5</sub>S | 0.5 µg/ml | 1 µg/ml | 1 µg/ml | 2 µg/ml | 0.5 µg/ml |

### Peptoid TM5 kills *P. aeruginosa* Xen41 faster than traditional antibiotics

Previously, *Pseudomonas aeruginosa* was tested against various peptoids to determine the bacterial killing kinetics ^49^. In the previous manuscript, TM1 is referred to as peptoid 1, TM5 is referred to as 1-C13_4mer_, and TM6 is referred to as 1-11_mer_. Xen5 and Xen41 come from different parental strains, so verifying that kinetics were the same was deemed necessary ^50^. To verify that the kinetics of Xen41 killing were similar to the kinetics of Xen5, *P. aeruginosa* Xen41 was grown in a similar manner to that described previously. Again, serial dilutions from 0.5 to 64 µg/mL were made and tested against Xen41. Each sample was done in triplicate and luminescence was measured at 0, 5, 10, 15, 20, 30, 45, 60, 90, and 120 minutes. TM5 showed a reduction of luminescence after 5 minutes, for all concentrations greater than 4 µg/mL (Figure 3a). After 45 minutes, there was near complete killing for concentrations of 16 µg/mL and above, while concentrations of 8 µg/mL appeared to inhibit growth (Figure 3a). In comparison, ciprofloxacin did not show reduction of overall luminescence until 45 minutes for all concentrations and did not show complete reduction for any concentrations for the entire 120-minute timeframe (Figure 3b). Interestingly, TM1 shows a decrease in luminescence for all concentrations after 5 minutes but shows a consistent decrease for concentrations of 8 µg/mL and higher (Figure S1a). After 30 minutes, a consistent decrease of luminescence was also observed at 4 µg/mL (Figure S1a). TM6 shows a decrease in concentrations of 16 µg/mL and higher starting at 5 minutes and shows some level of stagnation at 8 µg/mL (Figure S1b). The gold standard antibiotic ceftazidime did not show a reduction of luminescence for any concentration of drug tested during the 120-minute time frame (Figure S2a). Kanamycin showed a slight reduction from 60 minutes onward at 64 µg/mL while meropenem showed a decrease in luminescence after 60 minutes for most concentrations, albeit limited after the 60 minutes (Figure S2b,c). As such, all three peptoids showed a reduction in luminescence much faster than traditionally utilized antibiotics.

**Figure 3:**
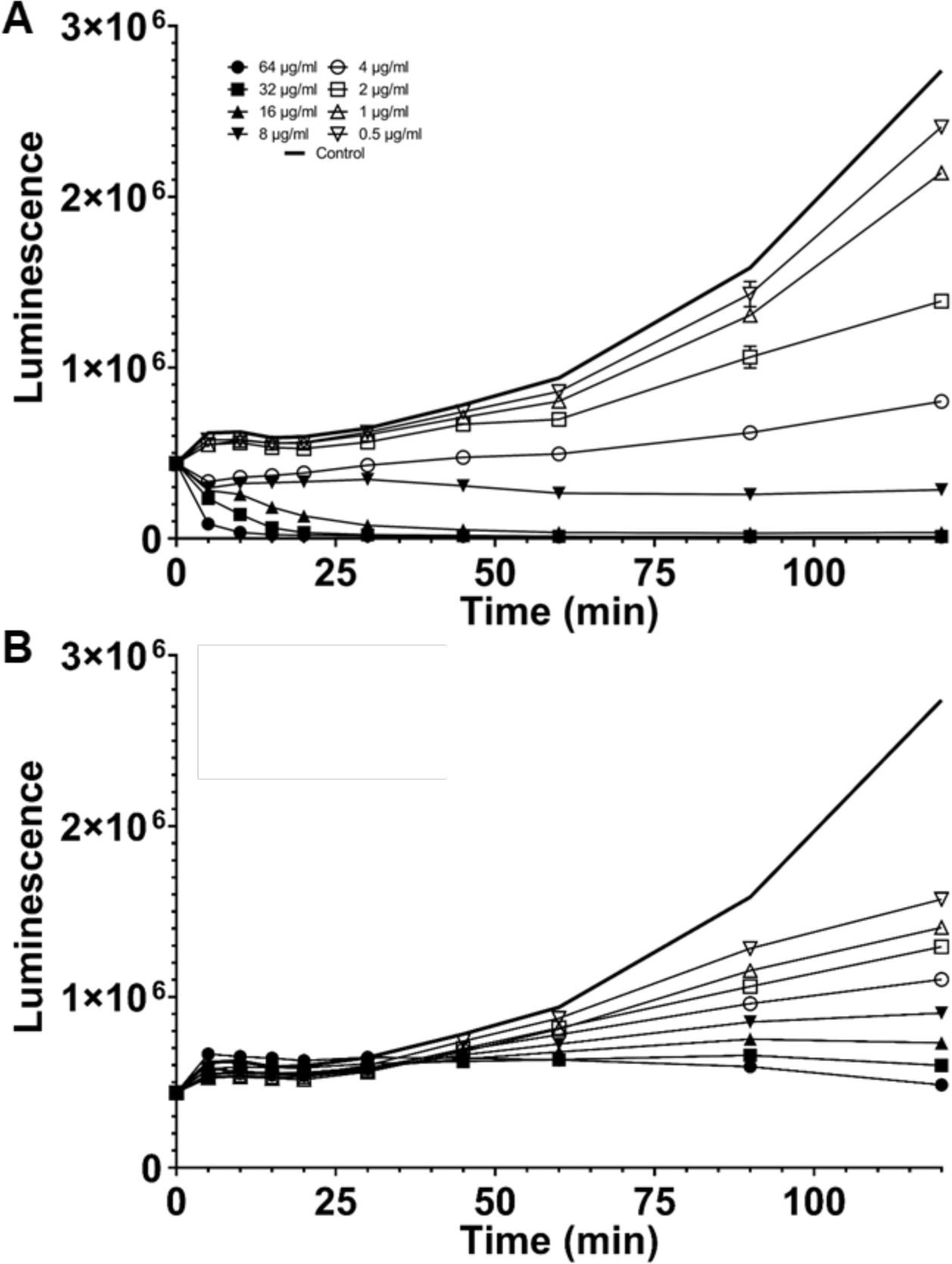
Time course experiments for 2-fold dilutions of TM5 (A) and ciprofloxacin (B) mixed with *P. aeruginosa* Xen41 were measured for luminescence at the following time points: 0, 5, 10, 15, 20, 30, 45, 60, 90, and 120 minutes. In addition to 2-fold dilutions, a control with PBS was also measured.

### TM5 treatment is well-tolerated by mammalian cells

To determine whether TM5 treatment would be safe and effective as a clinical therapeutic, we first evaluated the effects of treatment with peptoids on mammalian cells *in vitro.* We chose to focus on cell types relevant to *P. aeruginosa* respiratory infections, including the mouse fibroblast cell line L929, J774 mouse macrophages, and the human lung epithelial cell line A549. In comparison to previous generations of peptoids used as controls, TM5 was extremely well tolerated by A549 cells, showing no cytotoxicity in doses below 120 μg/ml using a MTC cell viability assay (Figure S3). We further tested the cells using and alamarBlue assay to examine whether the cells were actively producing energy and only saw mild effects on cell growth at concentrations above 32 µg/mL (Figure 4a). Using an MTS cell viability test, all mammalian cell lines tested showed no cytotoxicity observed in doses below 120 µg/mL, roughly 30-fold higher than the MIC of TM5 (Figure S3). This provided a large enough therapeutic window to justify further investigation of the effects of peptoids on mammalian cells.

**Figure 4:**
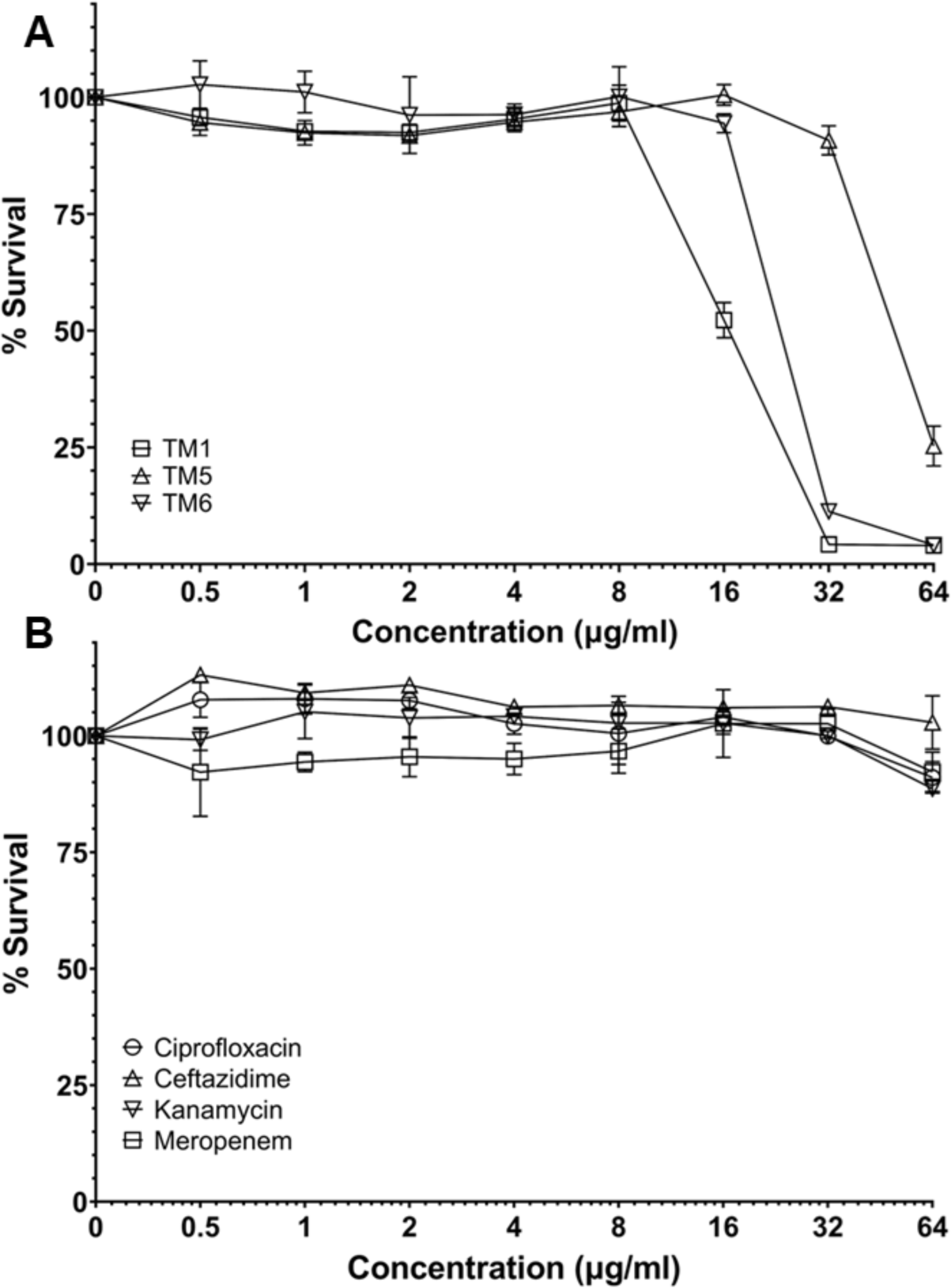
Cytotoxicity experiments with peptoids (A) and traditional antibiotics (B). An alamarBlue test was used to look at absorbance of 2-fold dilutions with *P. aeruginosa* Xen41. All concentrations and conditions were tested in triplicate.

### Development of tissue culture models of the mammalian lung for peptoid characterization

Although our initial data on the cytotoxicity of TM5 on mammalian cells was promising, the growth of tissue cultures cells in standard laboratory flasks and plates is not particularly well-representative of how these cells behave within a living organism. To better understand how the peptoid TM5 interacts with cells within the lung, we sought to develop more sophisticated tissue culture models that would allow epithelial cells to grow in conditions that promote these features.

We first developed and tested a spheroid model of A549 human lung epithelial cells to determine if the cytotoxicity of TM5 differed when cells were cultured with this method. Spheroids are simple organoids generally consisting of a single cell type grown as a scaffold-free three-dimensional model. Spheroids have previously been used extensively to study the delivery and toxicity of many novel cancer drugs as the pharmacokinetics and pharmacodynamics observed in these models more closely represent what is seen *in vivo* ^51–53^. More recently, they have also been used as models of viral pathogens as these multicellular clusters provide an ideal platform to observe viral growth and replication in a more complex microenvironment ^54^.

To form spheroids, A549 cells were grown in ultra-low attachment 96-well plates for 48 hours. To confirm the integrity of these spheroids, immunofluorescence analysis was performed by fixing the spheroids with 4% PFA and staining for actin (green) and the extracellular matrix protein laminin (red), and intact spheroids were observed (Figure 5a). The MTS assay was repeated with spheroid cultures, demonstrating that cells grown under these conditions show decreased cytotoxicity compared to flat cells, with only minimal cell death observed at a concentration of 240 µg/mL (Figure 5c).

**Figure 5:**
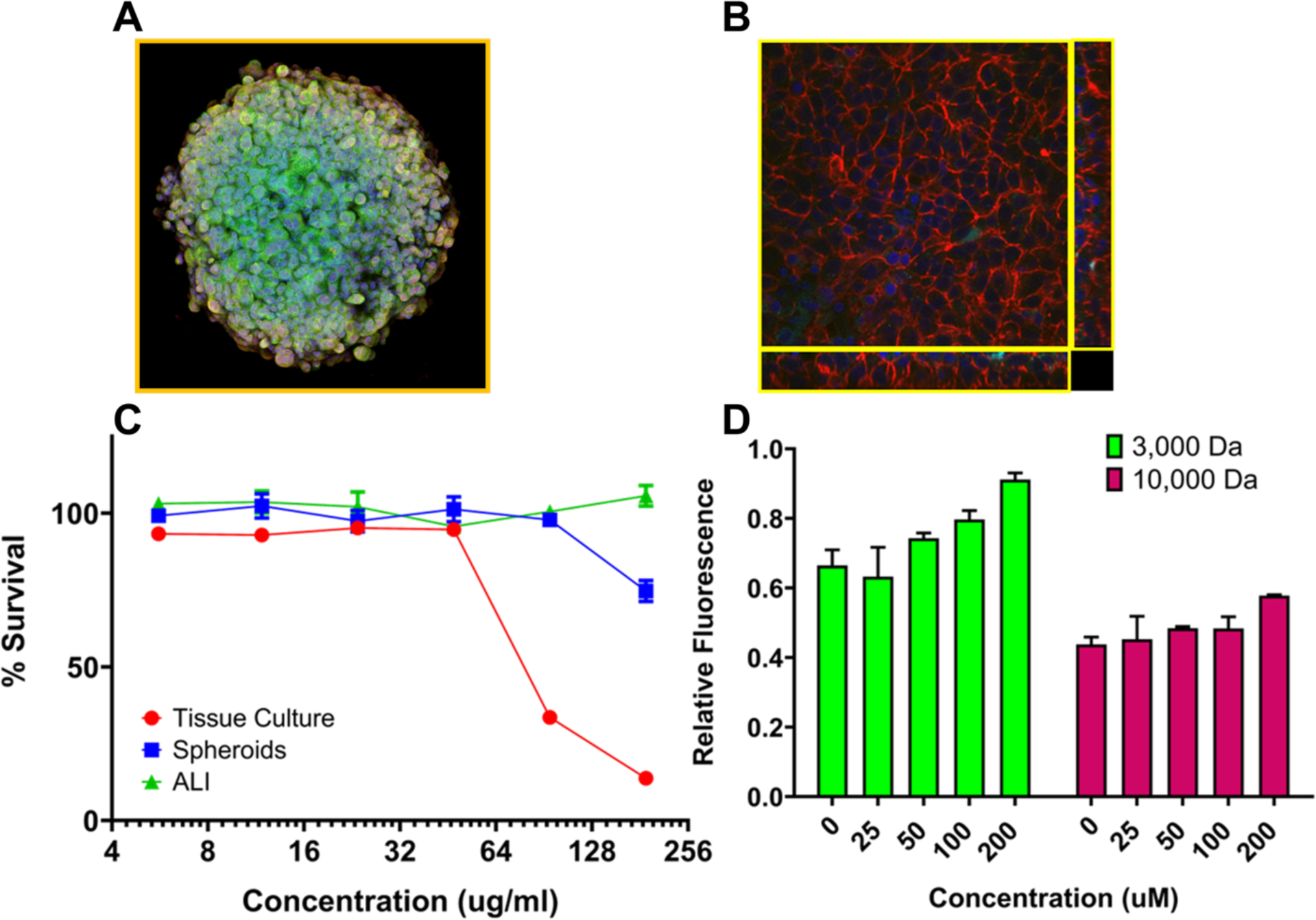
A549 cells were treated with peptoid for 3 hours, then analyzed by MTT assay. A549 cells were also grown in three-dimensional models either as (A) Spheroid cultures or (B) air-liquid interface (ALI) cultures. (C) Culturing A549 cells under conditions which mimic an in vivo lung environment result in decreased cytotoxicity following peptoid treatment (D) Integrity of the barrier function of ALI monolayers following peptoid treatment as measured by dextran exclusion assay showing that the monolayers remain intact following peptoid treatment. Dextran Alexa FluorTM 488 was used for the 3000 Da exclusion and Dextran Alexa FluorTM 647 was used for the 10,000 Da exclusion.

Next, we sought to demonstrate how epithelial cells would be affected by peptoid treatment in conditions where they are exposed to air to mimic the oxygen exchange environment in mammalian lungs using air-liquid interface (ALI) culture methods. Briefly, A549 cells were seeded onto transwell inserts and allowed to polarize for 10 days until fully confluent and displaying a cuboid phenotype. At this point, the media was removed to expose the confluent monolayer to air for an additional 48 hours. This exposure promotes differentiation of the epithelium, causing the cells to become ciliated, secrete increased extracellular matrix, and form more established tight junctions to create a tight barrier capable of excluding foreign materials. These ALI cultures were also fixed and stained for immunofluorescence with actin (red) and laminin (green) and analyzed by confocal microscopy. Z-stack images show the distinct cell junctions and thick cuboidal phenotypes characteristic of ALI cultures (Fig 5b).

When the same cytotoxicity assay was performed on ALI cultures, no cytotoxicity was observed at any concentration tested, suggesting that the differentiated cells were protected from any potential adverse effects when grown in conditions that best mimic the lung environment (Figure 5c). To further confirm this, we tested the barrier function of the ALI cultures using a dextran diffusion assay (Figure 5d). In this assay, fluorescently labelled dextran molecules of varying molecular weights are added to the apical surface of the culture. An intact and healthy barrier epithelium is able to exclude these sugars, preventing any of the dye from reaching the basal chamber beneath the monolayer. Even following three-hour peptoid treatment no significant decrease in barrier function was observed at any concentration of TM5, confirming that the A549 cells remained alive and healthy and able to perform their primary protective function.

### Intratracheal treatment with TM5 is safe and results in minimal adverse effects *in vivo*

Having established that TM5 causes extremely low cytotoxicity to mammalian cells cultured in three-dimensional models, we predicted it would demonstrate similarly low toxicity *in vivo.* Using a Balb/C mouse model, we initially inoculated a small number of animals with 50 µL of TM5 via the intratracheal route using doses that represented 5X and 10X the *in vitro* MIC for *P. aeruginosa* Xen41 (40 µg/mL and 80 µg/mL). These doses would be predicted to kill or inhibit bacterial growth without causing significant cytotoxicity to host cells. Indeed, we observed 100% survival of all animals following two rounds of treatment 24 hours apart, with no observable adverse effects on the animals’ health or well-being (Fig 6A).

**Figure 6:**
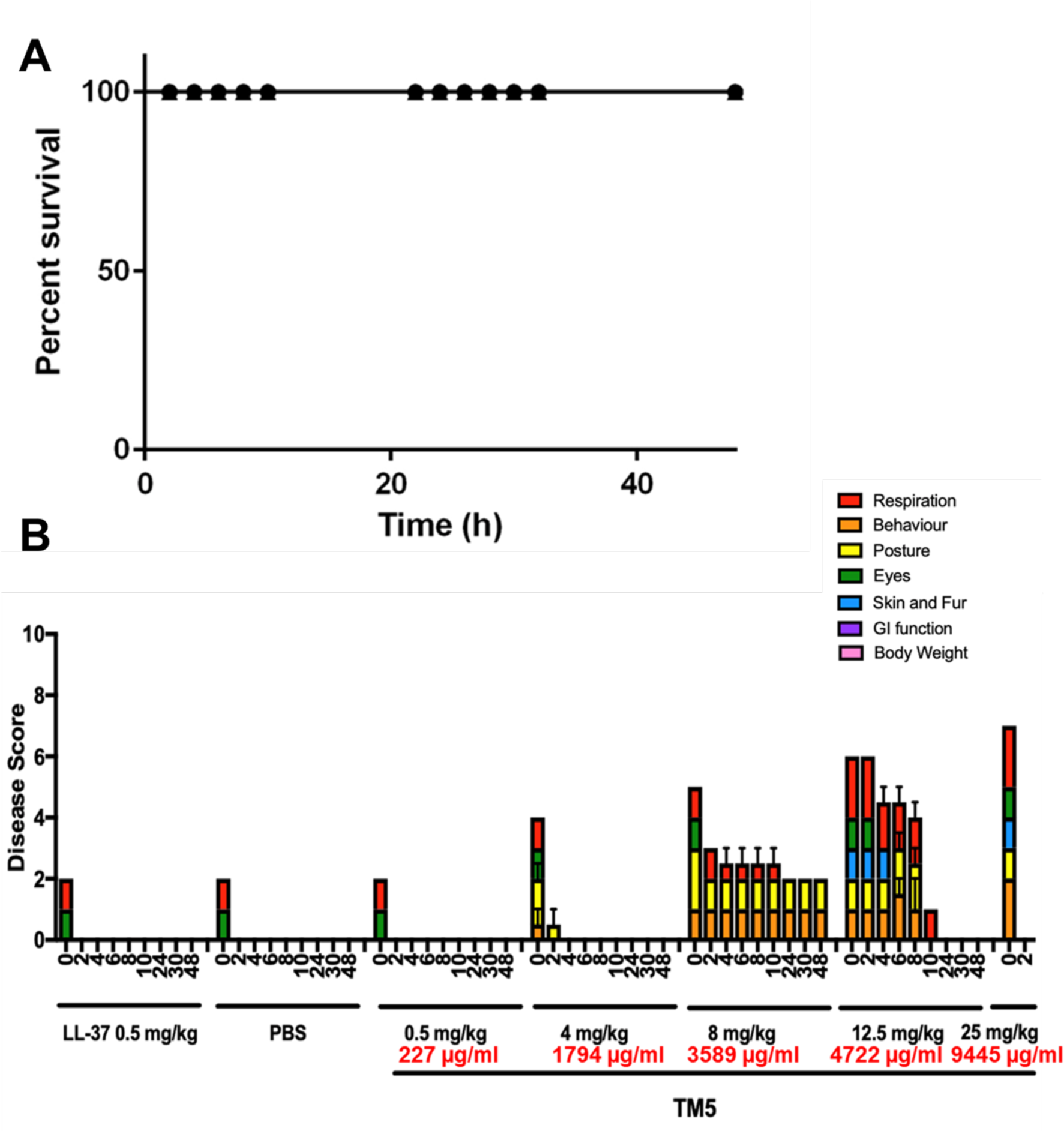
In vivo Toxicity of Peptoid TM5 following intratracheal inoculation (A) Balb/c mice were treated with 50 uL of either 40 ug/mL (•) or 80 ug/mL (▪) doses of TM5 and monitored for survival for 48 hours following treatment for signs of toxicity. (B) Balb/c mice were treated with increasing doses of TM5 to determine a toxicity threshold. No significant effects were observed at concentrations below 1.8 mg/mL

To determine the maximum safe dosage of TM5, we next treated animals with increasing intratracheal doses of the peptoid compared to a PBS control and the parental peptide LL-37. Animals were monitored every hour following treatment and scored using a modified Karnofsky score to measure their behavior, posture, weight, temperature, breathing, and physical appearance (Table S1). At approximately 40 µg/mL, our proposed therapeutic dosage, no effects above those seen in the controls were observed, and all animals appeared healthy and active within 30 minutes of treatment (Figure 6b). Even at concentrations 10-fold higher, minimal adverse effects were observed. To confirm the validity of our methods, we increased the concentration of antibiotics and noticed a gradual increase in toxicity reaching an LD_50_ at approximately 20 mg/mL, the maximum solubility of the peptoid. We therefore concluded that TM5 is safe for *in vivo* use at over 2500X MIC concentrations.

### Peptoid TM5 significantly reduces bacterial load in the lungs of infected mice compared to other peptoids

Since TM5 was well-tolerated when administered to the lungs in mice, the next step towards determining the efficacy of TM5 was testing it *in vivo* against *P. aeruginosa* infections in the lung. Groups of 8 Balb/c mice were each tested with TM1, TM5, TM6, and ciprofloxacin. 10 mice were set separated as untreated control mice and 4 mice were set aside as uninfected, untreated control mice. After administration of bacteria and treatment, 5X MIC, mice were monitored carefully, recording modified Karnofsky scores for each mouse over the course of the 3-day experiment (Figure 7). Untreated mice showed the highest disease scores on average after 24 hours and increased to 48 hours post-infection (Figure 7). Both TM1 and TM5 mice showed mild disease scores which decreased after 24 hours (Figure 7). Although TM6 had the highest average disease score of the peptoids, there was a large reduction after 48 hours (Figure 7). Mice treated with ciprofloxacin showed low overall disease scores and looked like the uninfected, untreated mice control after 48 hours (Figure 7). Based off disease scoring and the need for determination of CFUs after 24 hours, mice were sacrificed accordingly. Each day, one uninfected, untreated mouse was sacrificed as a control for imaging and CFUs. After 4 hours, two untreated mice were sacrificed to determine if the mice had received the proper number of bacteria. In addition, one mouse treated with TM6 met the criteria for sacrifice after 4 hours. After 24 hours, four mice from each group were sacrificed. One extra mouse from the TM6 group required sacrifice due to a high Karnofsky score. After 48 hours, three more untreated mice and one TM1 treated mouse required sacrifice due to high Karnofsky scores. After 72 hours, only one untreated mouse and two TM6 treated mice remained, while three TM1 treated, four TM5 treated, and 4 ciprofloxacin treated mice remained. All remaining mice were sacrificed accordingly.

**Figure 7:**
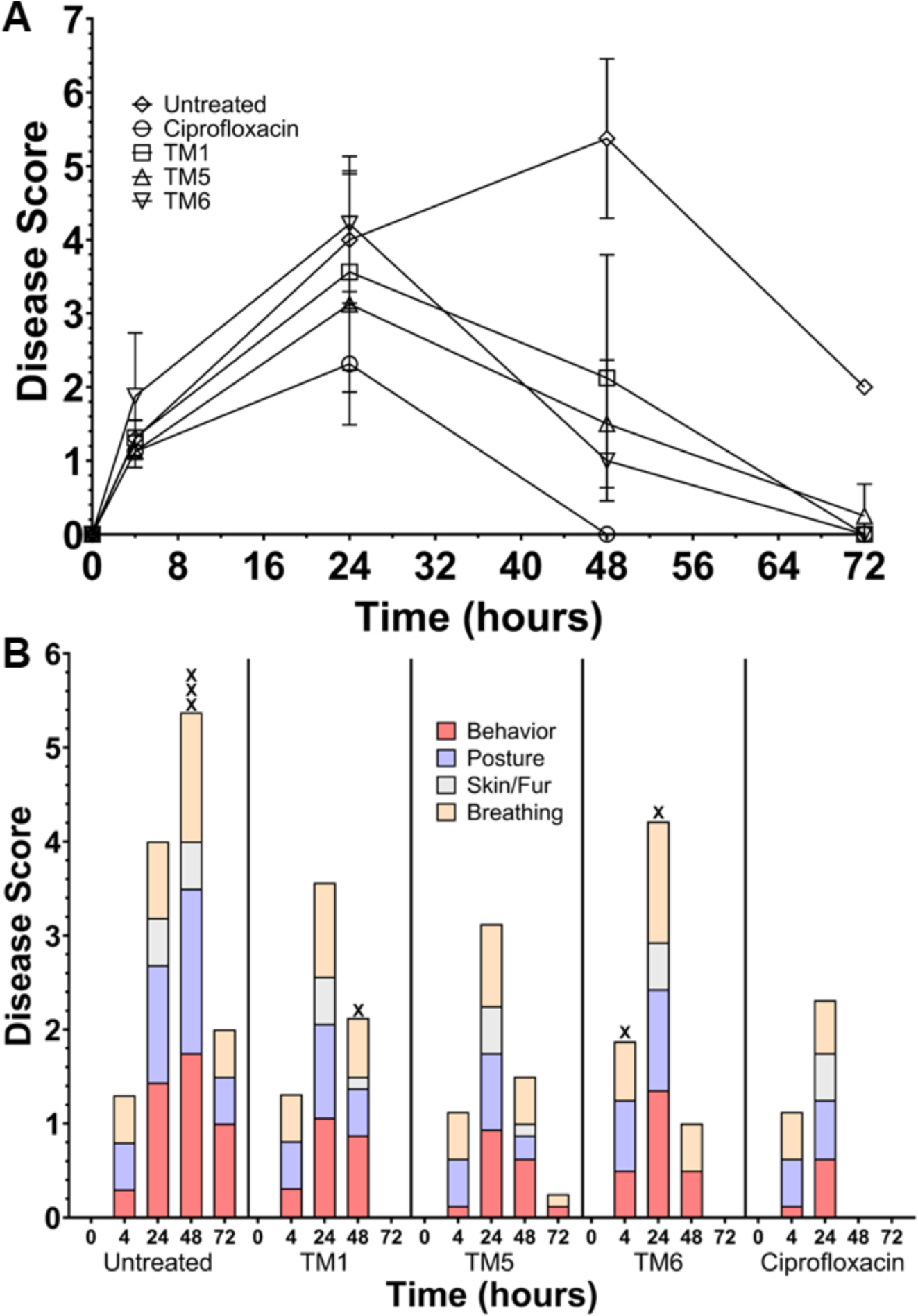
Representative Karnofsky scores. (A) Average overall Karnofsky scores for each group of mice tested from 0 hours to 72 hours. (B) Average Karnofsky scores for each group of mice over time, split into various categories tested. Columns marked with an ‘X’ indicate where a mouse was sacrificed ahead of schedule.

After sacrificing the mice, 1/4^th^ of each organ collected was homogenized in PBS and serially diluted to determine the bacterial load in the lungs and spleen of the mice. Untreated mice showed the highest average CFUs/mL in the lungs after one day of treatment, at about 2.2 x 10^7^ CFUs/mL (Figure 8a). Interestingly, the TM1 treatment group showed the next highest bacterial loads at 1.7 x 10^7^ CFUs/mL, followed by the TM6 treatment group at 1.4 x 10^7^ CFUs/mL (Figure 8a). Mice treated with TM5 showed significantly decreased bacterial CFUs in the lungs, at 4.1 x 10^6^ CFUs/mL, while ciprofloxacin showed the best overall CFUs in the lungs at 6.5 x 10^4^ CFUs/mL (Figure 8a). In both ciprofloxacin and TM5 treatment groups, one mouse was determined to be a significant outlier using the Grubb’s test for outliers with a p-value of 0.05. As a result, these data were removed for the lung CFUs. Similarly, one untreated mouse and one ciprofloxacin treated mouse were removed from the spleen dataset as significant outliers. Overall, mice treated with TM6 showed the highest CFUs in the lungs at 1.2 x 10^6^ CFUs/mL, followed by the untreated mice at 1.1 x 10^6^ CFUs/mL and the TM1 treated mice at 1 x 10^6^ CFUs/mL (Figure 8b). Peptoid TM5 showed the lowest CFUs/mL for the peptoids tested again, this time in the spleen, at 5 x 10^5^ CFUs/mL (Figure 8b). Ciprofloxacin showed the lowest CFUs/mL in the spleen at 2.9 x 10^3^ CFUs/mL (Figure 8b). It should be noted that for both the lungs and spleen, ciprofloxacin-treated mice were able to completely clear infection in two mice after one day for *P. aeruginosa* Xen41.

**Figure 8:**
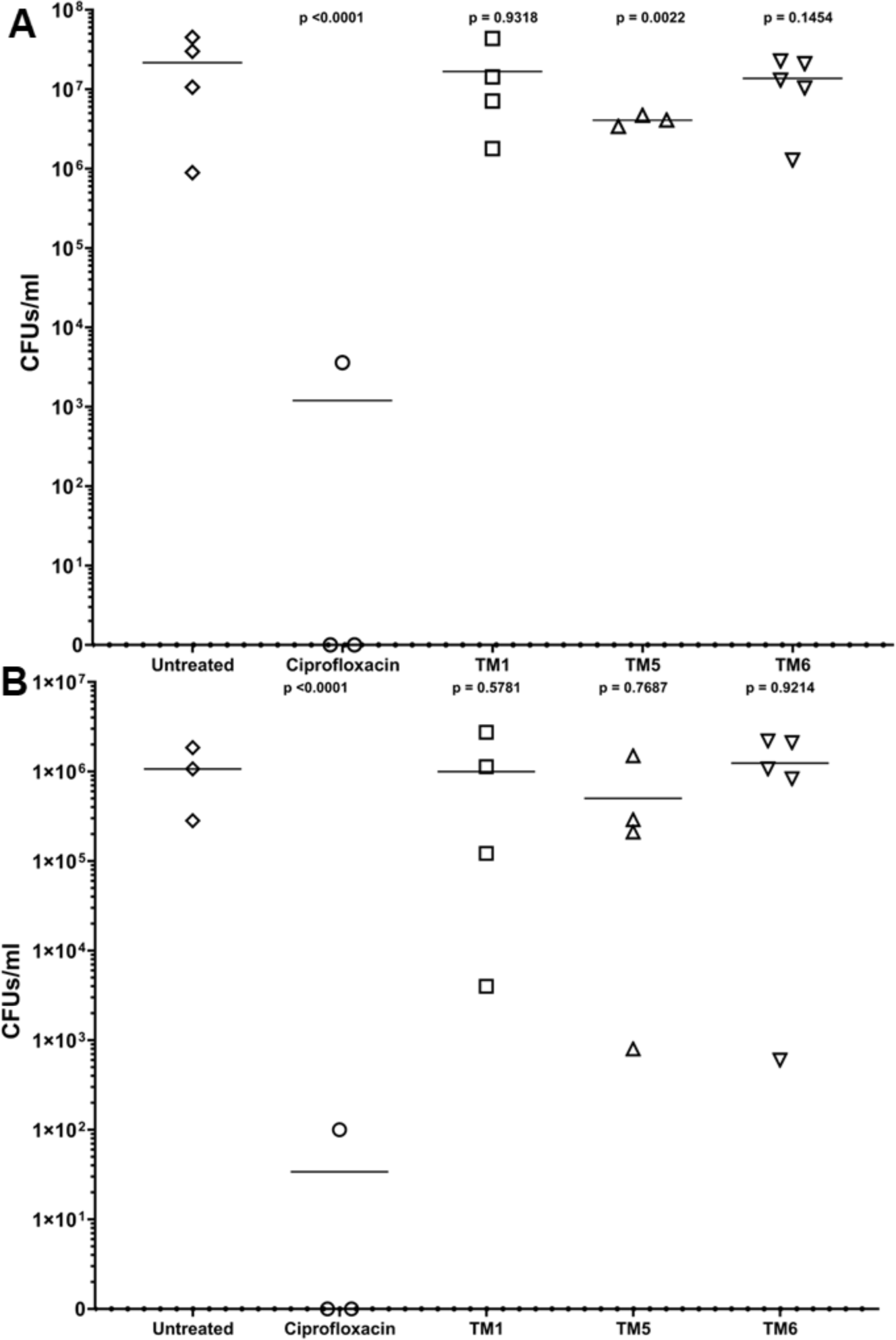
CFUs taken from sacrifice of day 1 mice. (A) CFUs/ml of Xen41 in the lungs of mice from varying treatment groups. (B) CFUs/ml of Xen41 in the spleen of mice from varying treatment groups. For all graphs, p-values are compared to the average of untreated mice. The line across represents the average of all data points within the group.

Throughout all *in vivo* experiments, mice were monitored via *in vivo* imaging using the IVIS system was used to measure luminescence within infected mice. Initial experiments showed readable levels of luminescence in mice 6 hours post-infection with 10^7^ CFUs of Xen41; however, survival rates of these mice were minimal after 24 hours (Figure S4). As a result, a lower dosage, 10^6^ CFUs of Xen41, was used to infect mice. This dosage was below the IVIS limit of detection, resulting in no measurable signal *in vivo*. However, necropsy of sacrificed animals and subsequent further imaging of the exposed organs revealed visible differences in all treatment groups compared to the untreated control (Figure 9).

**Figure 9:**
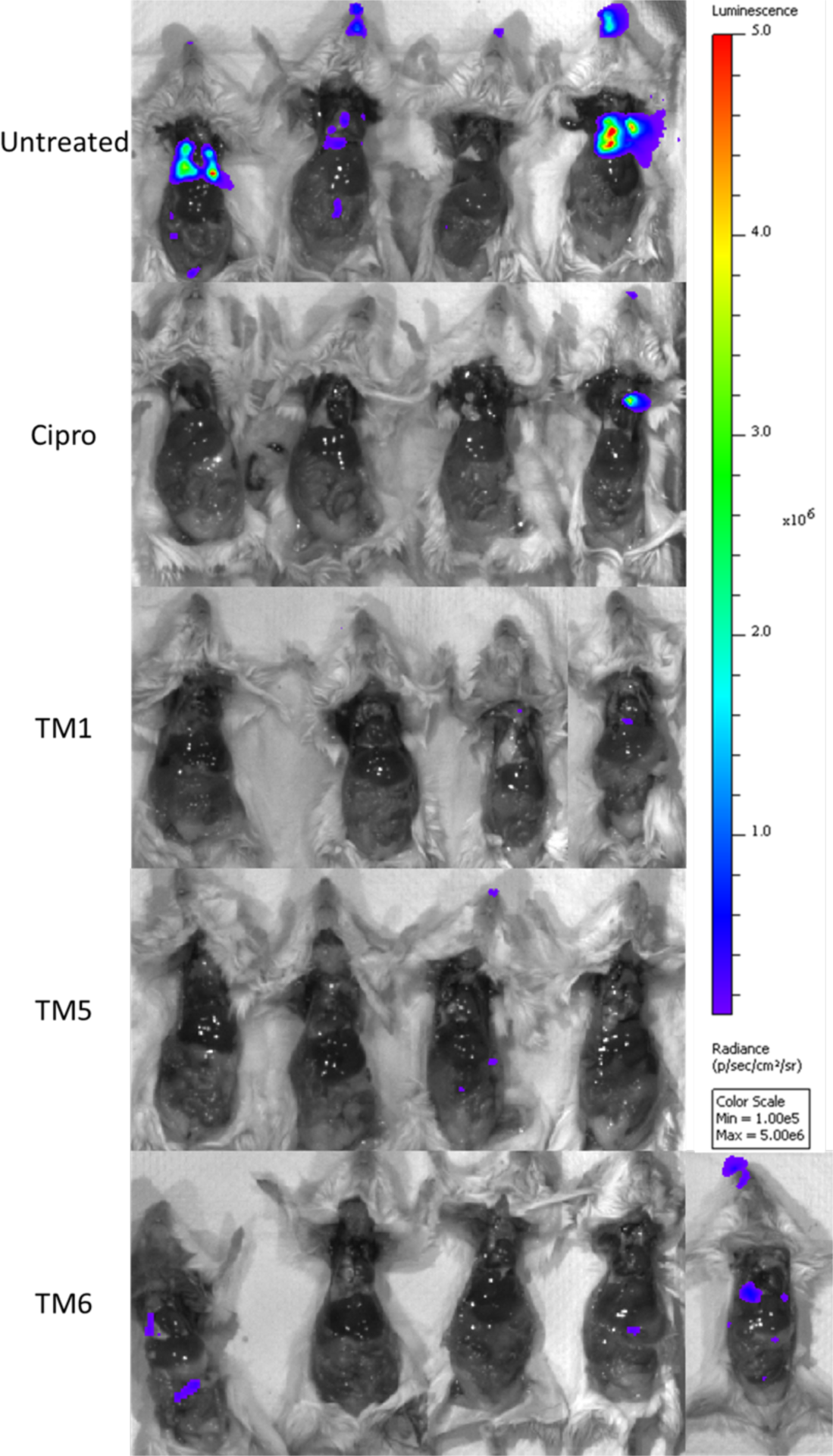
IVIS imaging of mice using the same scale of radiance for all mice tested. Each group of mice initially sacrificed were imaged together. An additional mouse in the TM6 group was imaged later using the same scale of radiance as it met the euthanization criteria.

### TM1 and TM5-treated mice demonstrated increased inflammation and neutrophil recruitment to the site of infection

Once CFUs had been determined, histology was performed to determine the effect both bacterial infection and treatment had on the lungs of the mice. Several main categories of effect were investigated such as, but not limited to, purulent inflammation, necrosis, and distribution. Scoring was compared between the individual mice and between the different treatment groups (Figure 10a,b). When the totals were averaged, mice treated with ciprofloxacin showed the lowest overall average score of 5.8. TM6 showed the next lowest score, 7.4, followed by untreated mice at 9.6. TM1 showed the second highest score at 13.6 and TM5 had the highest average score at 15.5. Looking at individual mice, all but one of the ciprofloxacin treated mice showed scores under 5 and all but one of the TM6 treated mice showed scores under 6. Untreated and TM1 treated mice showed relatively even distribution of scores between mice. TM5 treated mice showed the most consistency across mice, showing identical scores across all mice within the treatment group.

**Figure 10:**
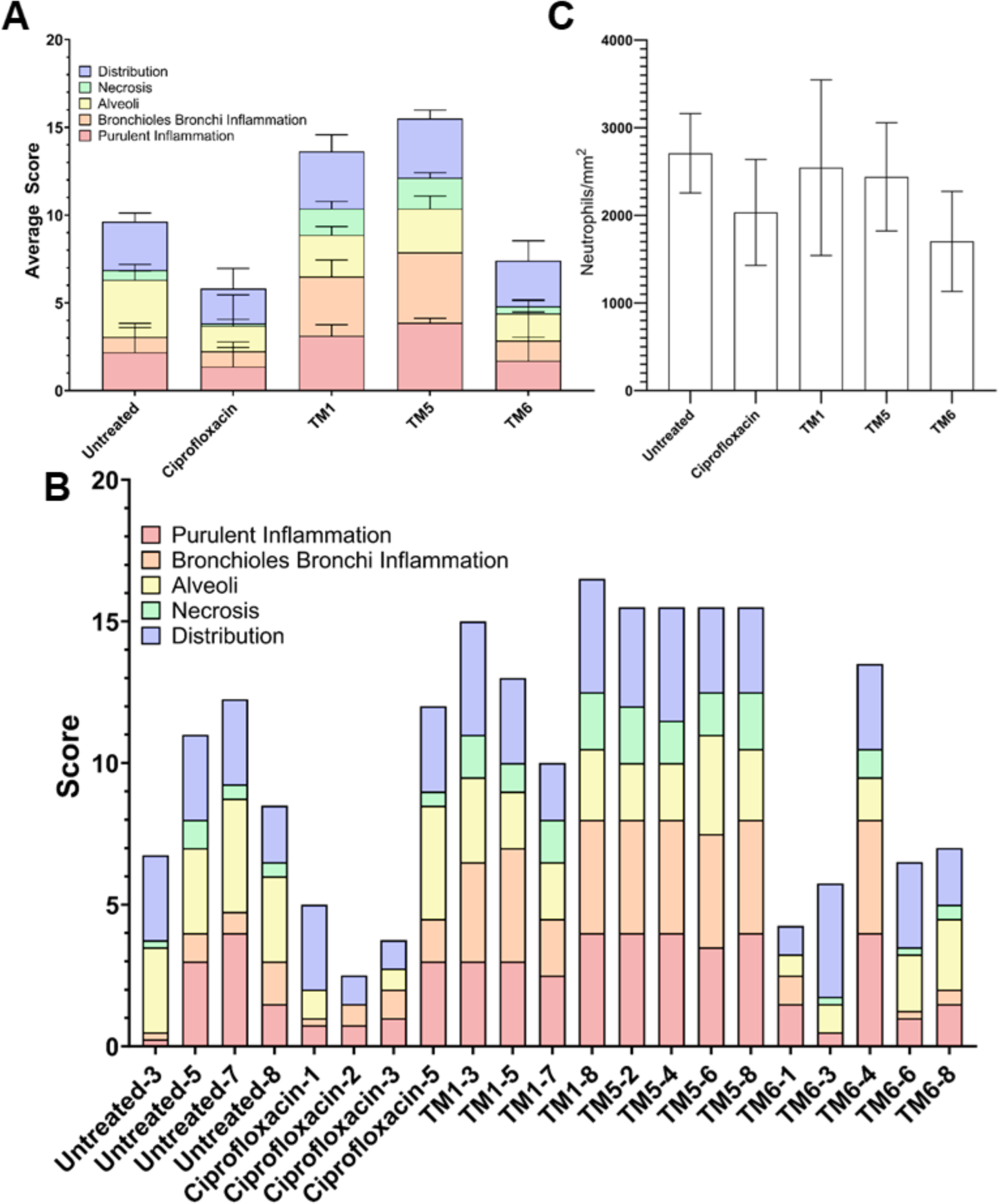
Breakdown of histology scoring as (A) averages of groups with standard error for each category and (B) individual scoring across several categories for each mouse sacrificed at day 1. (C) Neutrophil counts per mm^2^ for histology sections with standard error via computational analysis.

Nearly half of the total histology scoring for TM1 and TM5 mice appeared to be either purulent or bronchioles bronchi inflammation. As a result, computational determination of neutrophils per mm^2^ was determined (Figure 11c). Despite some variation between mice, ciprofloxacin and TM6 treated mice appeared to have lower levels of neutrophils per mm^2^, between 1800 and 2000, than untreated, TM1, or TM5 mice. Untreated, TM1, and TM5 treated mice all appeared to have relatively similar counts of neutrophils per mm^2^ at about 2500-2800. The correlation between increased inflammation and neutrophil recruitment in TM5 treated mice with improved disease scores and bacterial loads leads us to conclude that peptoids treatment leads to enhanced progression towards the recovery stage of disease. TM6, interestingly, showed the least amount of inflammation in the histology sectioning, while TM1 and TM5 showed high levels of inflammation in various parts of the lung sections (Figure 11). Ciprofloxacin treated mice also showed low levels of inflammation and untreated mice showed higher inflammation (Figure 11).

**Figure 11:**
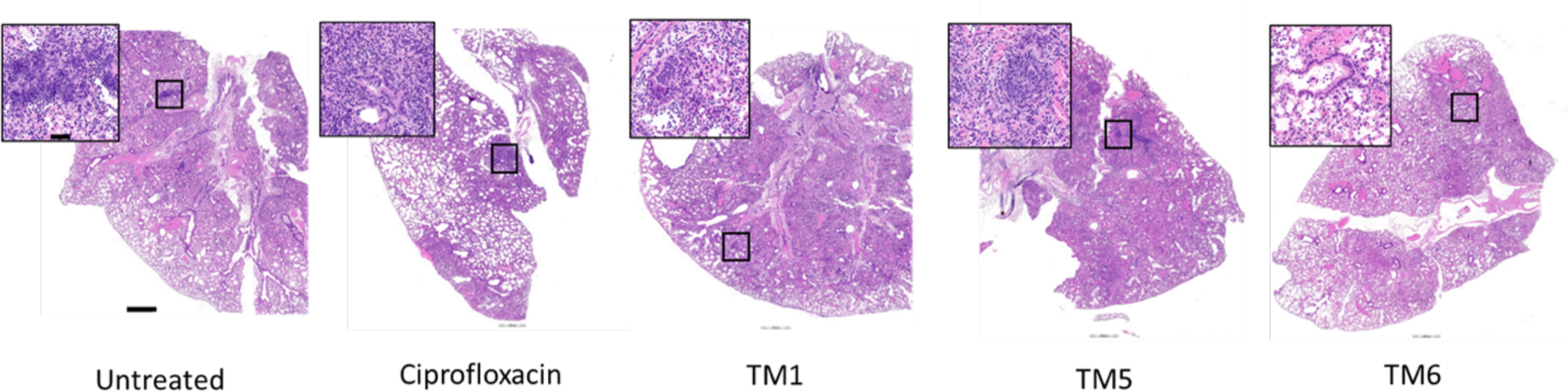
Histology sectioning of lungs for representative mice in each group. Insets are increased magnification of the box seen in the larger image. The short bar, in the inset, is equivalent to 50 μm and the longer bar is equivalent to 500 μm

## Discussion

The need for novel antimicrobials that are effective against MDR pathogens has become well-established as a research imperative. The work presented in this study represents an exciting avenue for reaching this goal by investigating a promising class of potential therapeutics and defining a potential treatment route and therapeutic window in which they are effective against *P. aeruginosa* respiratory infections. The peptidomimetics used in this work are a step forward in improving upon previous attempts at developing antimicrobial peptide-based drugs. While a small number of peptide-based antibiotics including polymyxin B, vancomycin, and daptomycin have been approved for clinical use, the development of natural AMPs and their derivatives as antimicrobials has not been easy. ^55^ Although a number of such drugs have gone through clinical trials, most have not reached clinical use due to their degradation and/or toxicity within a host.^56^ The peptoids described here have been specifically designed to counteract the problem of degradation by host proteases, and previous work has shown them to be extremely stable ^57, 58^ This work also addresses the question of toxicity within a host, with focus on a respiratory route of administration and treatment, demonstrating that peptoids are a viable treatment option for lung infections. Furthermore, we have demonstrated significant efficacy against *P. aeruginosa* respiratory colonization in an *in vivo* model, opening the door for development of these peptidomimetics for treatment of these serious infections.

Due to the promise that TM5 and other similar peptoids have shown previously ^37, 40, 42, 59^, it was necessary to utilize a strain of Pseudomonas that was both representative and allowed for proper MIC/MBC determination. As such, *P. aeruginosa* Xen41 was chosen as a bioluminescent strain produced from wild type *P. aeruginosa* PAO1. Bioluminescence is a good measure of bacteria viability due to the necessity of ATP production, so in addition to standard OD, MICs were also determined for this strain by comparing the bioluminescence. This technique provides additional unique information regarding treatment with sub-MIC concentrations of peptoids which suggests differences in mechanisms of inhibition in TM5 compared to the other peptoids. While all 3 peptoids showed a decrease in luminescence after 4 hours of incubation at concentrations above 1 µg/mL, TM5 shows a 5% larger decrease (Figure 1a). In addition, TM5 also showed a little over 75% reduction in luminescence at 2 µg/mL, while TM1 only saw that level of reduction between 4 and 8 µg/mL. Both TM1 and TM5 showed nearly complete reduction of luminescence at 8 µg/mL, while TM6 required concentrations of 32 µg/mL and higher to achieve the same reduction. This together suggests that TM5 was more effective at reducing the metabolic activity of Pseudomonas at lower concentrations.

Hydrophobicity and self-assembly have been shown to play important roles in the activity of peptoids. For example, a previous study showed that simply cutting the 12mer TM1 in half, yielding the 6mer TM7, resulted in little to no activity nor evidence of self-assembly in solution, while inclusion of a hydrophobic C13 alkyl tail to the design of the 5mer TM5, results in a very active compound since the tail boosts the overall hydrophobicity of the short peptoid ^36^. While the main repeating unit is the same in each of these three peptoids, an *N*lys-*N*spe-*N*spe unit, the *N*tridec tail in TM5 may allow it to better interact with the membrane at lower concentrations. Furthermore, due to the presence of the alkyl tail, TM5 self-assembles into ellipsoidal micelles, in contrast to the helical bundles formed by TM1 or TM6 ^36^. As such, it is likely that both hydrophobicity and self-assembly play larger role in the greater efficacy of TM5 over TM1 and TM6 in this study.

In terms of viability of Pseudomonas, TM1 treatment produced a significant reduction in CFUs/mL at 0.5 µg/mL and above, while TM5 showed a significant reduction in CFUs/mL at 2 µg/mL and above (Figure 1b). TM5 did, however, show a larger decrease in overall CFUs between 2-4 µg/mL compared to TM1, and both showed no viable bacteria at concentrations of 32 µg/mL and above. TM6 showed a steady decline of CFUs/mL, reaching a significant reduction at 4 µg/mL and no CFUs at 32 µg/mL. Much of this could be due to experimental variability, at least at concentrations of 4 µg/mL and above, but the overall MBC remained consistent. The data supports the idea that TM5 may be more efficient at lower concentrations than other peptoids, albeit similar at like concentrations towards the middle of the tested range. As such, this finding combined with our initial cytotoxicity data in cultured cells led us to test TM5 as our primary candidate against Pseudomonas, with the acknowledgement that TM1 showed promise as well.

To compare peptoid treatment with standard regimens, the traditional antibiotics ciprofloxacin, ceftazidime, meropenem, and kanamycin were also tested in this study since Xen41 has not yet demonstrated resistance to these drugs. Kanamycin showed the least change in bioluminescence when determining MICs, showing values of 32-64 µg/mL respectively (Table 2). As would be expected of frontline antibiotics, meropenem and ceftazidime were shown to have MIC_90_s of around 1 µg/mL, (Table 2). The MBCs for ceftazidime and meropenem were slightly higher at 2-4 µg/mL (Table 2). Kanamycin, on the other hand, showed an MBC of >64 µg/mL, consistent with its MIC. The difference in ceftazidime and meropenem MICs and MBCs could be explained by the observation that β-lactam antibiotics typically affect peptidoglycan biosynthesis and result in deformation, and subsequent collapse of the bacteria ^60–62^. Unlike these other traditionally used antibiotics, ciprofloxacin showed the greatest consistency across MIC and MBC, showing a significant reduction in both luminescence and CFUs/mL at concentrations of 0.5 µg/mL and higher (Figure 2). As a result, ciprofloxacin was utilized for all in vivo studies as the control antibiotic. Our results are consistent with previously determined MICs/MBCs ^63^ and make sense from the perspective of the mechanism of action ^64^. Since ciprofloxacin inhibits DNA gyrase, it is expected that bioluminescence would be low as well as the CFUs.

The time needed for most antibiotics to be effective depends inherently on the mechanism of action and the amount of bacteria present. In this study, it is apparent that antibiotics such as ciprofloxacin can act within one to two hours of incubation *in vitro* (Figure 3b). Interestingly, this is not the same for ceftazidime, meropenem, and kanamycin, all of which took more than two hours to observe a decrease in bioluminescence at all tested concentrations (Figure S2). When compared to the peptoids, it is clear that traditionally used antibiotics are measurably slower in terms of time-to-kill. For TM5, there is an immediate reduction of bioluminescence at 5 minutes for all concentrations at or above the MIC_90_ (Figure 3a). After 10 minutes, TM1 and TM6 show reductions of their bioluminescence at their MIC_90_s (Figure S1). When compared to the previous strain tested against these peptoids, Xen5, this time to kill is identical, suggesting that these peptoids are effective in the same time frame for various strains of *Pseudomonas aeruginosa* ^49^. This difference in time needed to kill the bacteria represents a strong advantage that peptoids have over traditional antibiotics.

The innate bactericidal properties of our antimicrobial peptoids demonstrated here in and previous studies ^37, 40, 42, 59^ suggests that they have promising potential as antimicrobial agents to help combat the escalating threat of AMR. However, to fully explore this potential it is important to thoroughly investigate any potential for off-target cytotoxicity against host cells, with the goal of defining a therapeutic window at which the peptoids can safely be used to treat clinical infections. Previous studies have demonstrated high specificity and relatively low cytotoxicity for all three peptoids explored in a number of cell lines ^40, 59, 65^. The effects of peptoids on lung cells had not yet been examined, and there are limitations to how accurately cell lines cultured in plasticware are able to represent the *in vivo* environment. As *P. aeruginosa* is of particular concern in lung infections associated with cystic fibrosis, we investigated the effect of TM1, TM5, and TM6 treatment on A549 human lung epithelial cells and compared these to L929 mouse fibroblasts by quantifying ATP production to determine the percentage of viable cells following peptoid treatment. We were able to demonstrate similar levels of cytotoxicity in these cells lines as has been previously shown (Figure 4, Figure S3), with peptoid TM5 showing the lowest cytotoxicity with little to no cell death observed at concentrations up to 64 μM.

Although cultured mammalian cells represent an excellent starting point for examining host toxicity, there has been a growing push towards developing more accurate models of human tissues to evaluate potential therapeutics, including for characterizing AMPs ^66–69^. *In vivo*, the epithelial cells that make up the lining of mammalian alveoli grow in a highly polarized fashion within the lung, with most of the respiratory epithelium consisting of pseudostratified columnar cells that function as a barrier between the airways and interstitial tissue. However, when cultured through traditional laboratory methods, lung epithelial cells grow with a flattened morphology in a single layer, do not show apical-basal polarity, do not form typical cell junctions, and fail to express extracellular matrix proteins. To more closely model the *in vivo* epithelium, complex 2 and 3-dimensional models such as organoids, organ-on-a-chip, and air-liquid interface cultures allow the examination of how molecules interact with cells in an environment that more accurately represents what occurs *in vivo*, and when human cells are used in such models, they avoid any potential species-specific effects that could undermine animal models. To assay our peptoids under more life-like conditions, we used two models constructed from A549 cells – spheroids and air-liquid interface (ALI) cultures. Spheroids have become popular in drug discovery as they demonstrate more accurate pharmacokinetics and pharmacodynamics compared to flat cells ^70, 71^, while ALI cultures are particularly valuable for studies of the human lung as they allow epithelial cells to differentiate into morphologies observed within the human alveoli ^72, 73^. We observed that in both models, A549 cells show greatly decreased cytotoxicity to peptoid treatment, demonstrating that higher concentrations of peptoids can be tolerated under conditions similar to those observed in the human lung (Figure 5). We hypothesize that this is likely due to the formation of tight junctions between cells which decreases the cell surface area exposed to the drug by preventing diffusion of the molecules between cells. For an extracellular pathogen such as *P. aeruginosa*, this would imply that while exposure to the bacteria would remain similar, the host cells would have decreased binding of the peptoids and thus lower side effects and toxicity. We believe these models are likely to represent how the peptoids would be encountered in patients, suggesting that these drugs will be well tolerated by lung epithelial tissues at significantly higher concentrations.

To confirm that peptoid TM5 would display similarly low toxicity levels *in vivo* as were observed in our *in vitro* lung epithelium models, we designed an intratracheal drug administration model in BALB/c mice. In this way, the peptoids could be applied directly to the lung tissues via intubation to determine if any negative effects were observed. For our initial study, we administered doses equivalent to 5X and 10X of the MIC for TM5 (20 μg/mL and 40 μg/mL respectively) and monitored the mice for any ill effects. No significant changes in behaviour or signs of illness were observed at these concentrations, and there was no impact on mouse survival. Following the success of this pilot study, we proceeded to administer increasing concentrations of TM5 to determine the threshold at which we would begin to observe adverse effects, which we monitored using a modified Karnovsky scale of disease severity (Table S1). In line with our *in vitro* data, we saw no change in behaviour, posture, skin and fur grooming, eye irritation, respiration, gastrointestinal issues, weight, or body temperature above what was observed for the PBS control at 400 μg/mL (100XMIC) concentrations. This is also similar to what was observed for the parental AMP, LL-37, which was administered as a control. LL-37 has been evaluated in human clinical trials and demonstrated to be safe by oral and topical routes,^74, 75^ so it is notable that TM5 shows a similar lack of toxicity through the intratracheal route.

As mice were dosed with increasing concentrations, we began to observe transitory effects on respiration and behaviour as would be consistent with minor irritation of the lung tissue at extremely high concentrations between 3200-10,000 μg/mL. These represent concentrations far beyond what would be used in a clinical setting, but nevertheless we observed that all these symptoms had fully resolved within 24-48 hours post-inoculation. This suggests that the peptoids are extremely well-tolerated by intratracheal inoculation and are likely to be safe to administer far beyond the recommended dose. In fact, no effects on mortality were observed until peptoids were administered at the solubility point of the peptoids, with an LD_50_ of 20,000 μg/mL or 25 mg/Kg observed. The fact that mortality was observed around the solubility point suggest that any ill effects may have been due to the peptoids precipitating out of solution. The tolerance of TM5 observed *in vivo* is notable because to the best of our knowledge it represents the first demonstration of a well-tolerated peptide-based treatment through intratracheal instillation. The development of novel therapeutics through this route has wide implications for the treatment of lung infections without risk of toxicity to other tissues or through systemic treatment and is likely to be particularly useful in *P. aeruginosa* therapeutics due to the predominance of lung infections with this pathogen in patients with cystic fibrosis.

Since TM5 showed little to no toxicity at much larger than 5x MIC levels, all three peptoids were tested alongside ciprofloxacin, to determine and compare their *in vivo* efficacy. A modified Karnofsky score, similar to that used in the *in vivo* cytotoxicity study was applied to monitor the mice. Each infected mouse received 10^6^ bacteria through a 20 µL intratracheal administration, followed by a 20 µL administration of treatment or saline control. Overall, mice treated with TM6 or saline demonstrated poor Karnofsky scores after 24 hours, while TM5 and TM1 showed moderate scores at that same time point (Figure 7). As would be expected for a frontline antibiotic, ciprofloxacin-treated mice demonstrated the lowest scores, although all mice showed some discomfort after 24 hours. All mice showed lower overall Karnofsky scores after 48 hours except the untreated mice and all treated mice appeared to show minimal symptoms after 72 hours post-treatment, suggesting that at this infectious dose, mice that can survive the initial onset of disease are eventually able to clear the infection.

In this time frame, 4 mice from each group were sacrificed at the 24-hour time point to observe the effect of treatment on the infection. During the timeframe of this experiment, several mice reached the euthanasia criteria of 3-4 points according to the Karnofsky score. Untreated mice were most severely affected to the point of requiring euthanasia, with only 1 mouse remaining at 48 hours post-infection. The TM6 treatment group had similar issues, resulting in one mouse being sacrificed after just 4 hours, and all but two mice sacrificed by the 48-hour timepoint. TM1 treated mice fared better, resulting in 3 mice remaining after 48-hour time points, while both TM5 and ciprofloxacin-treated mice had 4 mice remaining at the same timepoint. To better explain these results, imaging was performed throughout the experiment. However, for *P. aeruginosa* intratracheal infections, we observed an incompatibility between the limit of detection of *in vivo* imaging experiments and the infectious dose at which at least 50% of the animals are able to survive the infection (LD_50_). In preliminary experiments, a bacterial burden of 10^7^ CFU was utilized so that luminescence could be readily quantitated via the IVIS system (Figure S4). Unfortunately, this dosage resulted in nearly all animals requiring euthanasia, including those which were treated. As such, a lower dosage, 10^6^ CFUs, was utilized for all future experiments. The log fold decrease resulted in the majority of mice surviving the course of the disease and even clearing the infection after 3 days; however, the mice exhibited no measurable external bioluminescence. However, after sacrificing mice and imaging them *ex vivo*, some luminescence was observed in the lungs and spleen, confirming the hypothesis that luminescence was occurring just below the limit of *in vivo* detection. (Figure 9).

The disease scores and *ex vivo* imaging data observed in all groups correlated well with bacterial burdens as measured when looking at CFUs. Ciprofloxacin treated mice appeared to have near baseline levels of *P. aeruginosa* in the lungs after 24 hours (Figure 8a). Similarly, the TM5 group showed significantly reduced CFUs in the lungs compared to the untreated mice. TM1 and TM6 treated mice did not show any significant reduction in bacterial burden in the lungs, which could explain why they appeared sicklier than TM5 and ciprofloxacin-treated mice. Due to the speed at which these peptoids can eliminate bacteria, we predicted that a decent portion of the higher Karnofsky scoring was due to the cytotoxic effects of having large amounts of recently killed bacteria in the lungs. As such, histology was necessary to see the effect that infection and subsequent treatment had on the lungs of these mice.

Histology scoring showed that ciprofloxacin treated mice had less overall inflammation and necrosis on average when compared to all other treatments. Interestingly, both TM1 and TM5 appear to have the highest values overall, with nearly 50% of the scoring from inflammation (Figure 10a,b). This, in conjunction with the time-kill data of these peptoids, suggests that both TM1 and TM5 may have killed large amounts of the initial load of bacteria received. This would result in large amounts of potentially toxic debris to be left behind, which would likely recruit many of the inflammatory responses seen during infection. This is supported the higher levels of neutrophils localized in the lungs, similar to that of untreated mice (Figure 10c). As such, these data along with the lower number of CFUs in the lungs in TM5 treated mice suggest that mice were in the later stages of infection, possibly even into the beginning stages of recovery. Alternatively, as the peptoids are derived from LL-37, a natural product of the host immune system that can be conditionally pro-inflammatory and modulate neutrophil recruitment and response to infection, it is possible that TM1 and TM5 produce an immunostimulatory effect that enhances disease recovery ^28, 76^. H&E staining of sections of the lung seem to suggest that TM6 and ciprofloxacin may have immunosuppressing properties while TM1 and TM5 have immunostimulatory effects (Figure 11). A complete study is necessary to discriminate between these potential hypotheses and examine any potential effects of our peptoids on neutrophil function, however.

In this study, we have demonstrated the efficacy of TM5, an antimicrobial peptoid, against *P. aeruginosa* Xen41 and several clinical strains, both *in vitro* and *in vivo*. We also show the speed of bactericidal action is similar to that of previous studies, with killing occurring in a matter of minutes ^40, 49^. In addition, this study showed that the use of air-liquid interfaces (ALI) or spheroids to test cytotoxicity showed higher tolerated values compared to traditional culturing methods, like those seen in mice. The use of peptoids like TM5 has shown great promise, and as such, may be an alternative source for antimicrobials in the fight against resistant strains of various bacteria. Future studies looking at the various inflammatory markers and immunochemical parameters should be assessed to look at alternative uses for peptoids as some seem to promote the immune response while others seem to act as immunosuppressants. These studies would allow for a comprehensive strategy to better understand the mechanisms of peptoids *in vivo* and ultimately design the synthesis of new, more effective peptoids in the future.

## Materials and Methods

### Peptoid Design and Synthesis

Peptoid synthesis was performed as previously described ^38, 40, 77^. Briefly, peptoid synthesis was carried out using a Symphony X (Gyros Protein Technologies, Tucson, AZ) peptide synthesizer located at the Molecular Foundry in the Lawrence Berkeley National Laboratory, Berkeley, CA. Peptoids were synthesized on a Rink amide MBHA resin (EMD Biosciences, Gibbstown, NJ). All reagents were purchased from Sigma Aldrich (St. Louis, MO). Synthesis followed the submonomer protocol from Zuckermann, et al ^77^. Peptoids were cleaved from the resin by treating with trifluoroacetic acid (TFA):triisopropylsilane:water (95:2.5:2.5 volume ratio) for 10 minutes. A C18 column in a reversed-phase high performance liquid chromatography (HPLC) system (Waters Corporation, Milford, MA) was used for purification with a linear acetonitrile and water gradient with a compound purity greater than 95% as measured by analytical reverse-phased HPLC. Confirmation of the peptoid synthesis was determined using electrospray ionization mass spectrometry.

### Bacterial Strains and Culture

Bioluminescent Xen 41 Pseudomonas aeruginosa, derived from the parental strain PAO1 (obtained from Xenogen Corp. now part of PerkinElmer, Waltham, MA) was used for all experiments unless otherwise noted. For comparative efficacy experiments, the additional strains PA14, PAZK2019, PAZK69, PAZK3095, PAZK2870 from the Kolter collection (Oxford University) were used in this study. All strains were streaked onto either cation-adjusted Mueller-Hinton (MH) agar plates or in Luria-Bertani (LB) plates and grown at 37 °C. For each experiment use a single colony was grown overnight in MH or LB broth at 37 °C with shaking at 250 rpm.

### MIC and MBC Assays

To determine the minimal inhibitory concentration (MIC) of the chosen antibiotics ceftazidime, ciprofloxacin, kanamycin, and meropenem, and all peptoids a 96 well plate was setup with triplicates of a 2-fold dilution of the to be tested antimicrobial in MH broth. For antibiotic testing, a polystyrene plate was used while polypropylene plates were used for all peptoid experiments. Overnight cultures of each P. aeruginosa strain were diluted 1:100 and grown to log phase (OD_600_ between 0.4 – 0.9). Log phase cultures were diluted to OD_600_ 0.001 and then added to the plate. The plate containing 2-fold dilution of the antimicrobial and the bacterial strain was incubated overnight at 37 °C. Bacteria and MH broth served as positive controls and just MH broth served as a negative control. The next day, the wells were gently resuspended luminescence and/or OD_600_ was measured using a plate reader. Unless otherwise noted, MIC Is defined by the 95% decrease of the untreated bacteria. Experiments using Xen41 had bioluminescence measured at 4 hours post incubation with treatment way to look at actively growing bacteria.

To evaluate MBC values, a 10-fold dilution of each sample from the MIC 96 well plate was prepared. The triplicates were plated onto MH agar plates and incubated overnight at 37°C. The colony forming units (CFUs) were enumerated the next day.

### Time-Kill Assays

To determine how quickly antibiotics and peptoids were able to kill Xen41, a 96 well plate was setup with triplicates of 2-fold dilutions of all tested antimicrobials in MH broth. Log phase cultures were diluted to an OD_600_ of 0.01 before 100 μL of bacteria was added to each well of the plate. Samples were incubated at 37 °C and bioluminescence was measured at the following time points: 0-, 5-, 10-, 15-, 20-, 30-, 45-, 60-, 90-, and 120-minutes post-addition of bacteria.

### Cytotoxicity Assays

Cytotoxicity in mammalian cells was measured using a MTS assay as has been previously described ^40^. Briefly, A549 human lung epithelial cells (ATCC) were cultured in F12K media (Gibco) supplemented with 10% FBS. For traditional assays, 10^4^ cells were seeded into 96-well plates and allowed to adhere for 18 hours at which point media was removed and replaced with 2-fold dilutions of peptide diluted in PBS. Cultures were incubated at 37 °C for 3 hours, at which point 20 μL of CellTiter 96 Aqueous Non-Radioactive cell proliferation assay (Promega) was added to each well. After 2 hours of further incubation, the OD490 absorbance was measured using a plate reader. For spheroid cultures, assays were performed as above except that 10^4^ A549 cells were grown in Costar Ultra-Low attachment 96-well plates (Corning) for 72 hours at which point spheroids had formed. For ALI cultures, 10^4^ A549 cells were inoculated into Transwell chambers (Corning) and allowed to polarize for 10 days before the apical medial was removed for a further two days before applying the peptoids. All experiments were performed in triplicate, and to confirm the validity of the MTS assay were repeated using an alamarBlue assay (Thermo Fisher Scientific).

### *In Vivo* Toxicity Studies

Balb/c mice were treated with increasing dilutions of peptide suspended in 50 μL PBS via the intratracheal route to evaluate toxicity on the animals using an Endotracheal Intubation Kit (Kent Scientific). Following treatment mice were monitored for survival and scored on a scale of 0-16 for their physical and behavioral responses to peptoid treatment using the modified Karnofsky score defined below. To control for effects of intubation and liquid administration, control mice were inoculated with either PBS or the parental peptide LL-37 (Anaspec).

### *In Vivo* Efficacy Assays

46 Balb/c mice were purchased and acclimated for 1 week prior to any experiments in cages of 4, except for two cages of 5 mice. Once acclimated, mice were anesthetized using 2-3% isoflurane with oxygen. All subsequent imaging was carried out using the same anesthetization conditions. Mice were split into treatment groups consisting of 8 total mice per group except for untreated mice which consisted of 10 total mice, and uninfected mice, which consisted of 4 total mice. After anesthetization, mice were given a dosage of 10^6^ bacteria, administered intratracheally using a catheter in 20 μL aliquot. A gentle puff of air was used to ensure bacteria had been disseminated to the lungs before administration of 20 μL of treatment. Untreated mice were given 20 μL of saline to avoid differences in total volume given to the lungs between mice. Mice were then imaged for bioluminescence using an IVIS imaging system. Mice were closely monitored for the next 24 hours, with Karnofsky scoring done at 4-, 24-, 48-, and 72-hours post-infection. Mice which exceeded 4 points on the modified Karnofsky table were sacrificed immediately.

Initially, 1 uninfected and 2 untreated mice were sacrificed to get imaging controls as well as baseline numbers of bacteria that had been administered to the lungs. At 24 hours, 4 additional mice from each group were randomly selected to be sacrificed. The remaining mice were only sacrificed if meeting the Karnofsky scoring criteria as mentioned above until 72 hours, where all remaining mice were sacrificed. For all sacrificed mice, imaging was done after opening the mice to expose the lungs and other vital organs using the IVIS system, measuring for bioluminescence. After initial imaging, the lungs, spleen, and liver were harvested and imaged again. Lungs and spleen were then split for histology (stored in 10% neutral buffered formalin), CFUs (stored in PBS), and DNA/RNA isolation (stored in DNA/RNA Shield) and liver was collected for histology (stored in 10% neutral buffered formalin).

Lungs and spleen collected for CFUs were homogenized in PBS and collected for CFU determination. Each sample was serially diluted in 10-fold dilutions (n=3), plated on LB plates, and incubated at 37 °C overnight. Colonies which formed were counted and CFUs were back calculated.

### Histopathology

Following euthanasia, representative samples of lung were collected from each mouse and fixed by immersion in 10% buffered formalin for 48 hours at room temperature. Post-fixation, the tissues were stored in 70% ethanol before embedding in paraffin, sectioned at 5-6 µm, and stained with hematoxylin and eosin to evaluate tissue responses. The whole glass slides of histologic sections were scanned at 20x magnification using the Pannoramic Scan II by 3DHistec (Budapest, Hungary), and analyzed, in a blinded manner, by a board-certified pathologist, and ordinally scored for magnitude of tissue injury, cellular infiltration, inflammation and lesion distribution. Post-acquisition images were uploaded to Aiforia image processing platform (Aiforia Inc., Cambridge, MA) to quantify neutrophils via with a validated artificial intelligence/deep learning convolutional neural networks (CNNs) and supervised learning (Bencosme-Cuevas, Emily, Tae Kwon Kim, Thu-Thuy Nguyen, Jacquie Berry, Jianrong Li, Leslie Garry Adams, Lindsey A. Smith, Syeda Areeha Batool, Daniel R. Swa, Stefan H.E. Kaufmann, Yava Jones-Hall, Albert Mulenga. 2023. Ixodes scapularis nymph saliva protein blocks host inflammation and complement-mediated killing of Lyme disease agent, Borrelia burgdorferi. Front. Cell. Infect. Microbiol., IN PRESS). Statistical analysis of the histopathology ordinal data and neutrophils/mm2 was performed by using GraphPad Prism 10.

### Statistical Analysis

Statistical analysis was done using 2-way ANOVA, via GraphPad Prism, using multiple comparisons across different antibiotics and different concentrations of peptoid or antibiotic used.

## Acknowledgements

AEB thanks the NIH for funding this work with a Director’s Pioneer Award, grant # 1DP1 OD029517. AEB also acknowledges funding from Stanford University’s Discovery Innovation Fund, the Cisco University Research Program Fund, and the Silicon Valley Community Foundation, and Dr. James J. Truchard and the Truchard Foundation. We gratefully acknowledge Dr. Michael Connolly and Dr. Behzad Rad at the Molecular Foundry for assistance with peptoid synthesis and sample preparation equipment. Work at the Molecular Foundry was supported by the Office of Science, Office of Basic Energy Sciences, of the U.S. Department of Energy under Contract No. DE-AC02-05CH11231. MM thanks and acknowledges that portions of this work were supported by the National Institutes of Health under Ruth L. Kirschstein National Research Service Award # 1F32AI120589-01A1, and further thanks the University of Edinburgh Chancellor’s Fellowship and the UKRI MRC Precision Medicine Studentship funding MB.

---

**Figure S1:**
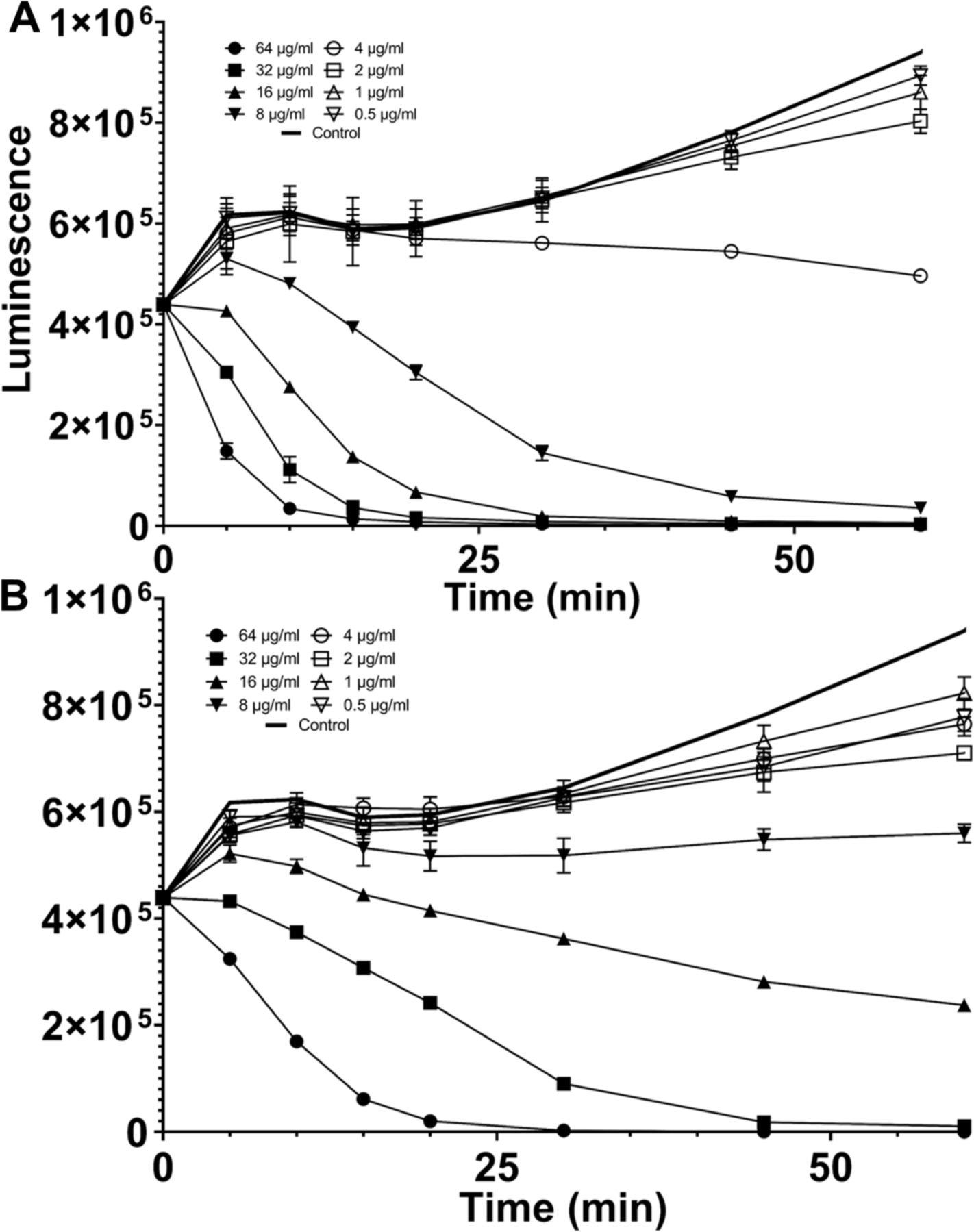
Time course experiments for 2-fold dilutions of TM1 (A) and TM6 (B) mixed with *P. aeruginosa* Xen41 were measured for luminescence at the following time points: 0, 5, 10, 15, 20, 30, 45, 60, 90, and 120 minutes. In addition to 2-fold dilutions, a control with PBS was also measured.

**Figure S2:**
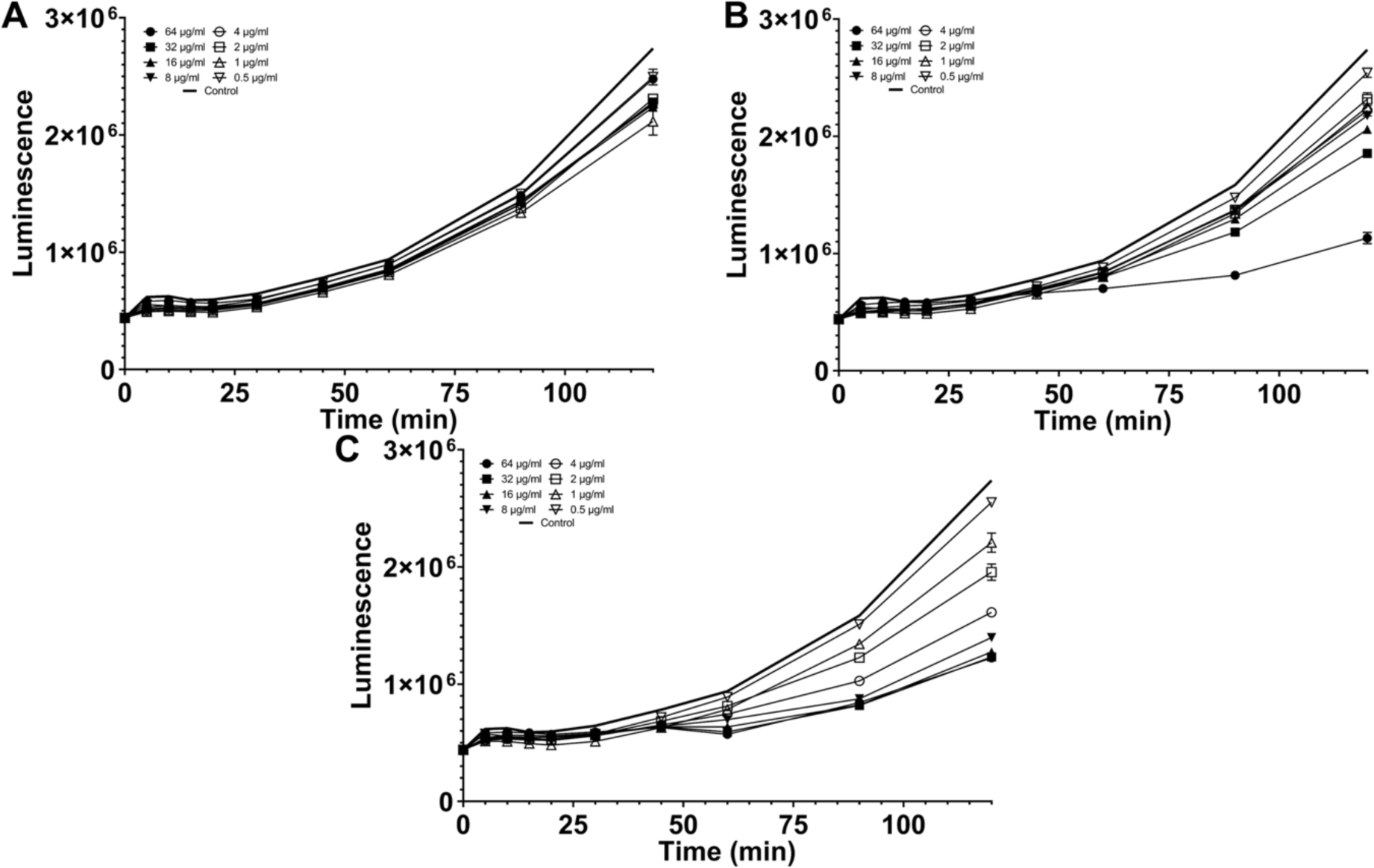
Time course experiments for 2-fold dilutions of ceftazidime (A), kanamycin (B), and meropenem (C) mixed with *P. aeruginosa* Xen41 were measured for luminescence at the following time points: 0, 5, 10, 15, 20, 30, 45, 60, 90, and 120 minutes. In addition to 2-fold dilutions, a control with PBS was also measured.

**Figure S3:**
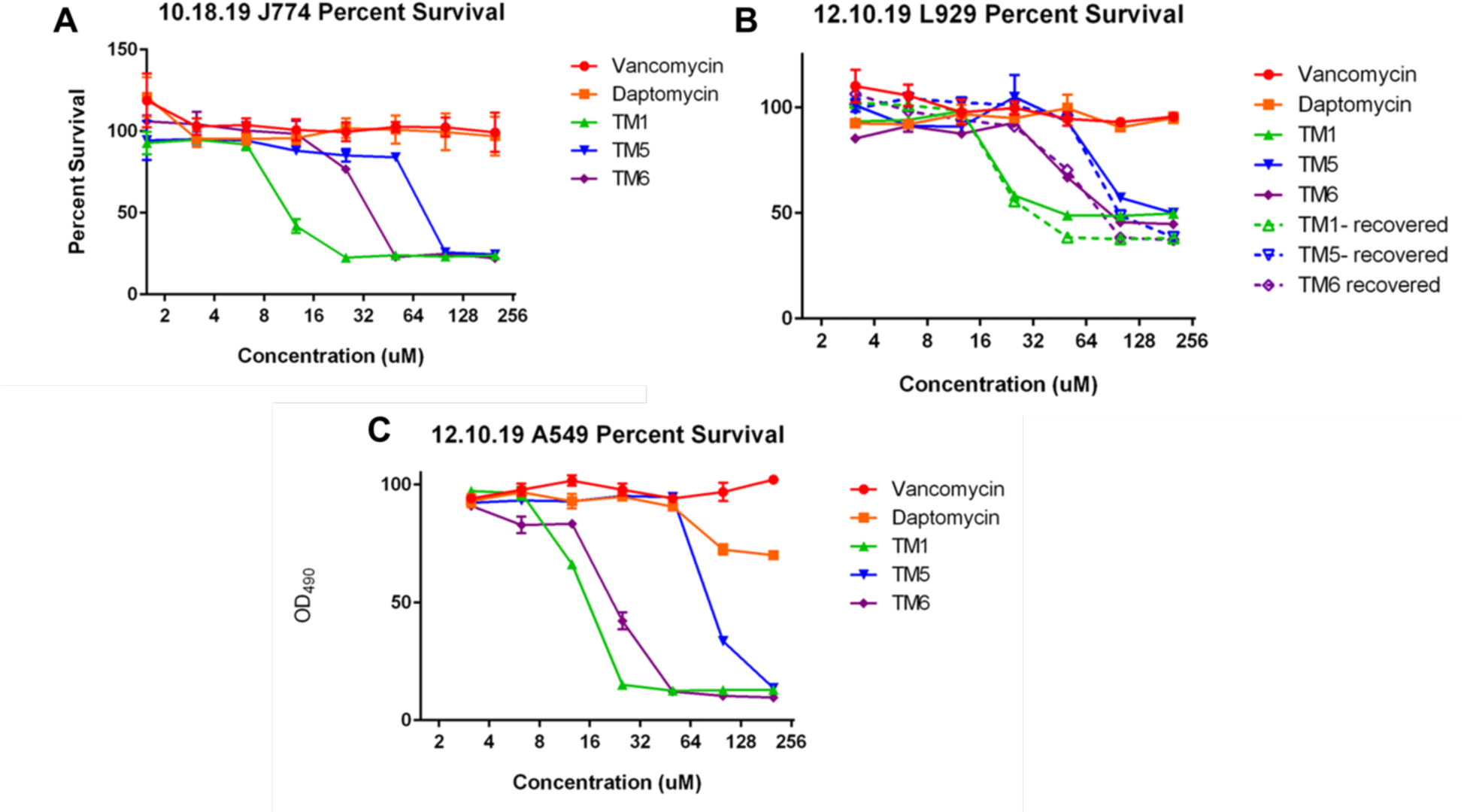
Cytotoxicity of TM5 and other related peptoids in mammalian cells. Cells were treated with peptoid for 3 hours, then analyzed by MTT assay to determine percent survival in (A) J774 mouse macrophages, (B) L929 mouse fibroblasts, and (C) A549 human lung epithelial cells.

**Figure S4:**
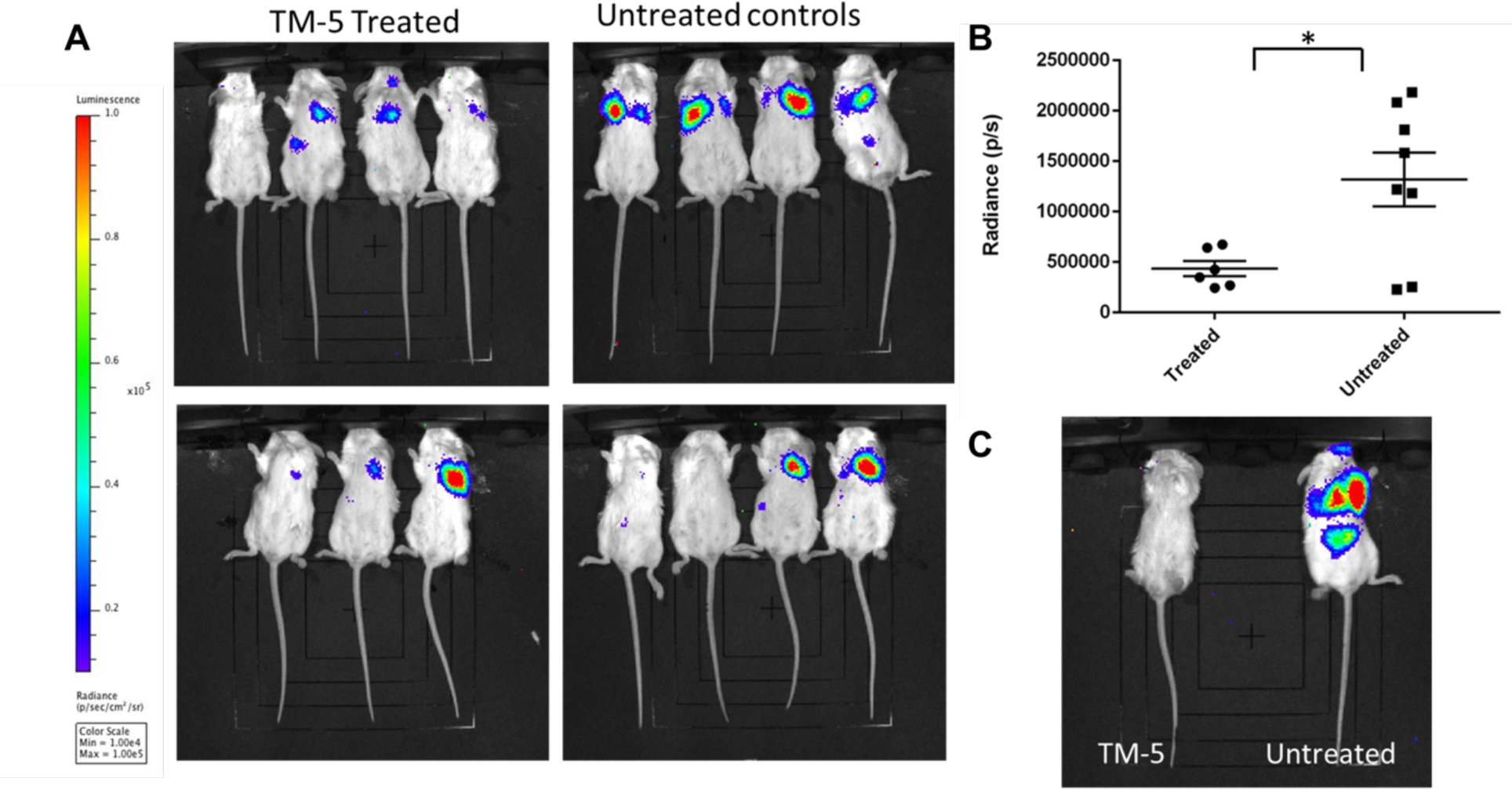
*in vivo* treatment of *P. aeruginosa* Xen41 infections with TM5 (A) *in vivo* imaging of TM5 treated (left) and untreated control (right) Balb/c mice 6 hours post-infection with 10^7^ CFU of *P. aeruginosa X*en41 via an intratracheal route. (B) Relative luminescence of animals at 6 hours post-infection (p= 0.0167) (C) *in vivo* imaging of surviving animals at 24h post-infection

**Table S1.** Modified Karnofsky Scoring of *in* vivo peptoid cytotoxicity. Animals were scored on a scale of 0-16 for their physical and behavioral responses to peptoid treatment following intratracheal inoculation.

| Scoring guide | 0 | 1 | 2 |
| --- | --- | --- | --- |
| **Behavior** | Alert, active, highly responsive | Reduced activity, slow responses | Lethargic, no response to stimulation |
| **Posture** | Normal | Occasional hunched posture | Hunched posture, unstable movement |
| **Skin and Fur** | Smooth, shiny, groomed | Mild ruffling, reduced grooming | Ruffled fur, no grooming |
| **Eyes** | Normal | Open w/ discharge or inflammation | Mostly closed, discharge and inflammation |
| **Breathing** | Normal | Nasal discharge | Forced breathing and nasal discharge |
| **GI tract** | Normal feces | Soft feces | Diarrhea |
| **Body temp** | 35-39°C (95-102°F) | 33-35°C(91-95°F) or 39-40°C(102-104°F) | <33°C or >39°C (<91°F- or >104°F) |
| **Weight** | Stable | 5-10% decrease | >10% decrease |

## Notes

### Competing Interest Statement

A.E.B. is a shareholder and member of the Board of Directors of Maxwell Biosciences, Inc., which is developing the antimicrobial peptoids for clinical use. No employee of Maxwell Biosciences had any role in the design of the study; in the collection, analyses, or interpretation of data; in the writing of the manuscript, or in the decision to publish the results.

### Summary of Updates

Author information updated

